# The Cellular Expression and Genetics of Purple Body (*Pb*) in *Poecilia reticulata*, and its Interactions with Asian Blau (*Ab*) and Blond (*bb*) under Reflected and Transmitted Light

**DOI:** 10.1101/121285

**Authors:** Alan S. Bias, Richard D. Squire

## Abstract

Mature Purple Body and Non-Purple Body male guppies differ from each other in several ways. Non-Purple males may have large numbers of xanthophores, erythrophores, and blue iridophores, in addition to the usual dendritic, corolla and punctate melanophores. Fewer violet iridophores are found. In contrast, homozygous Purple Body males lack collected and clustered xanthophores, although isolated single xanthophores remain. Violet iridophores and blue iridophores (violet-blue chromatophores units) abound. The dendrites of dendritic melanophores are finer and form chains with each other. Punctate and corolla melanophores in areas comprising orange ornaments are greatly reduced in number. The heterozygous Purple Body male has erythrophores similar to those of non-Purple males, but yellow pigment is reduced. The melanophores are not as greatly changed in orange ornaments. In Domestic Guppy strains, and at least in one suspected instance in wild-type, melanophore structure and populations may be further modified by one or more additional autosomal genes.

## Introduction

The intent of this paper is multifold; 1. To identify phenotypic and microscopic characteristics of newly described Purple Body trait **(Fig 1 and 2)**. 2. Provide photographic and microscopic exhibits of Purple Body and non-Purple Body for ease in identification of chromatophores types **(Fig 3A-E)** and their interactions. 3. To encourage future study interest at a cellular level of populations in which Purple Body highlights near-UV (Ultra-Violet) reflective qualities. 4. To stimulate future molecular level studies of Purple Body to identify linkage groups (LGs) which correspond to haploid number of individual chromosome(s) within the Guppy genome

**Fig 1.**
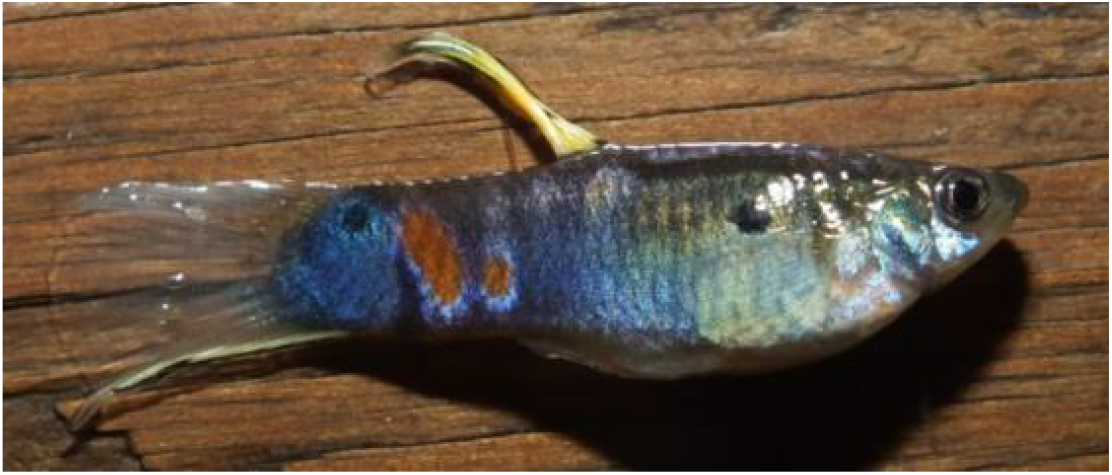
Homozygous *Pb/Pb male*

**Fig 2.**
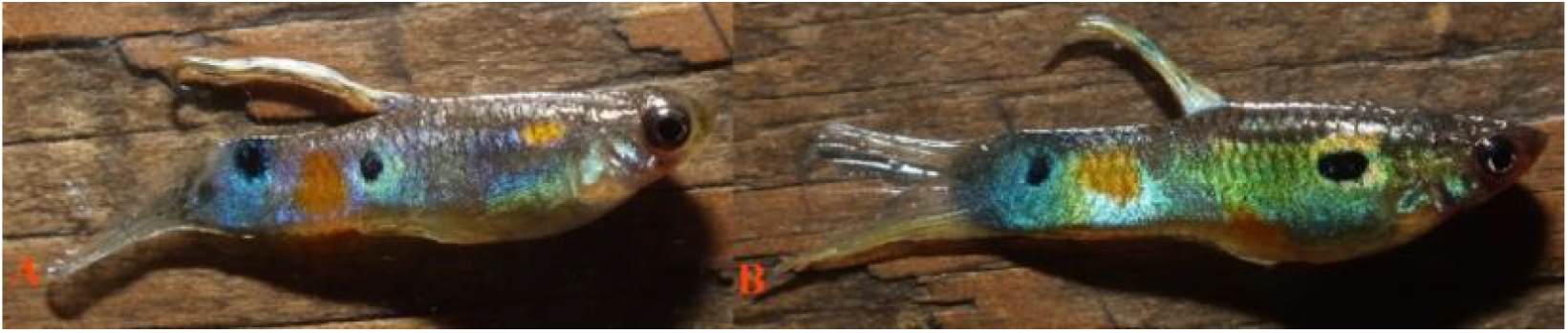
**(A)** Heteorzygous male *Pb/pb*, **(B)** Non-Pb *pb/pb* male

**Fig 3.**
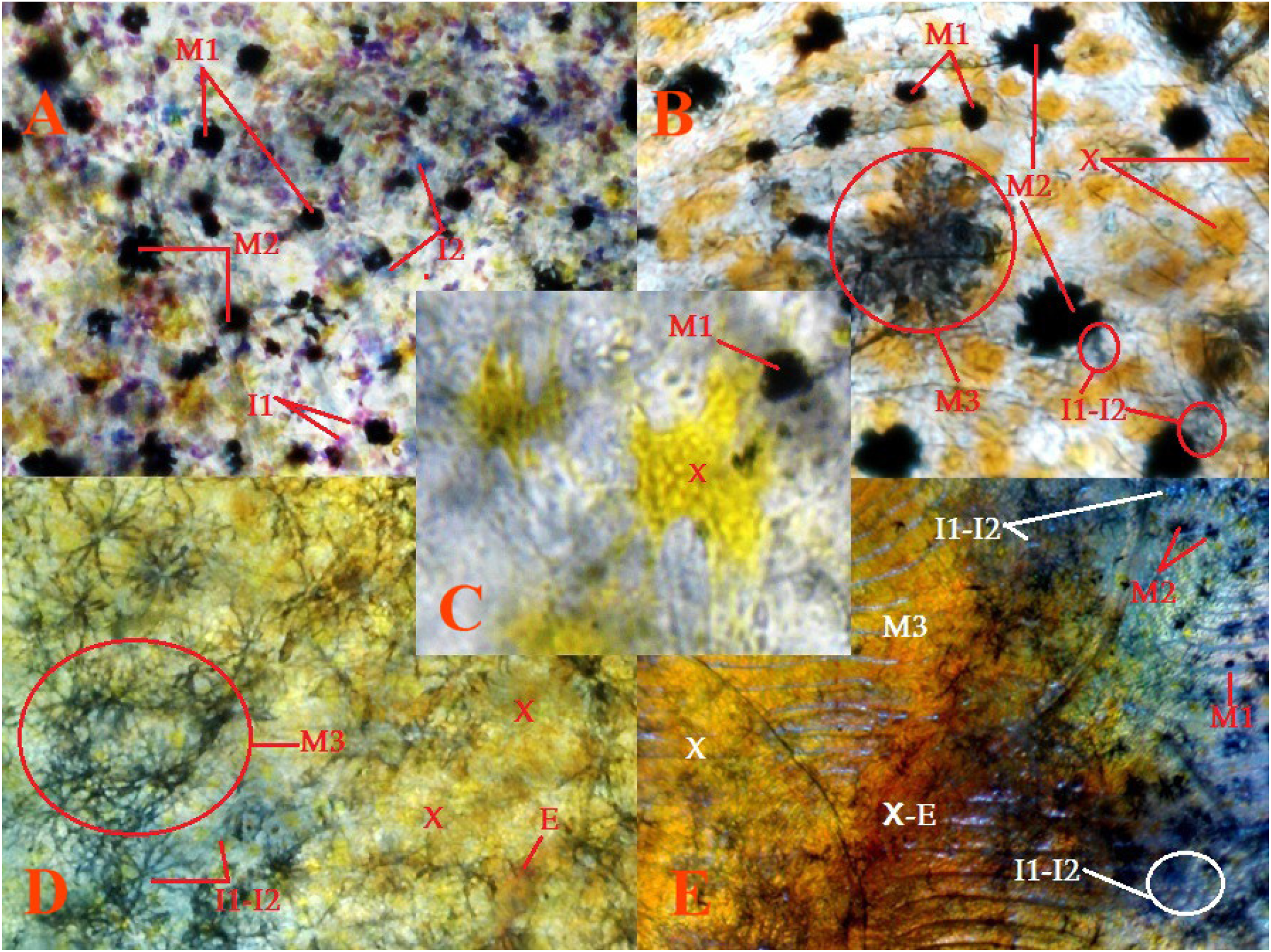
**Pigment cell types and structures identified, (A)** 100X, **(B)** 100X, **(C)** 100X, **(D) 40X**, **(E)** 40X. Melanophores Punctate (M1), Melanophores Corolla (M2), Melanophores Dendritic (M3); to include visible dendritic melanophore strings and violet/blue iridophore chromatophore units. Violet iridophores (I1), Blue Iridophores (I2); to include violet-blue iridophore collections. Erythrophores (E). Xanthophores (X); comprised of isolated single cells, collected cells, clustered groups (dendritic structures), dense collections or Xantho-Erythrophores (X-E).

## Materials

### ID Number, Pb or non-Pb, Color / Strain, Genotype

(***See:*** Supplemental S1 for Strain Genotypes and Slide Specimen Photos).

2 non-Pb [Reticulata Female] (grey E) *pb/pb*.

2 Pb [Top] male (grey E × F) *Pb/pb* and 2 non-Pb [Lower] male (grey E × F) *pb/pb*.

3 non-Pb male (grey E × F) *pb/pb*.

3 Pb [Reticulata Female] (blond E) *Pb/-*.

4 non-Pb Ab male (grey E) *pb/pb Ab/ab*.

5 Pb Ab male (grey E) *Pb/Pb Ab/ab*.

5 non-Pb male (blond) *pb/pb*.

6 non-Pb male (grey E × F) *pb/pb*.

6 Pb male (grey E × F) *Pb/pb*.

7 non-Pb male (grey E × F) *pb/pb*.

7 Pb male (blond) *Pb/pb*.

8 Pb male (grey E) *Pb/pb*.

9 Pb male (grey E × F) *Pb/pb*. Note: dried sample photo with constricted pigments, see 2 Pb for accurate color comparison.

13 Pb male (grey E) *Pb/Pb*.

14 non-Pb male (grey E × F) *pb/pb*.

15 Pb male (grey L - Pingtung) *Pb/pb*.

16 non-Pb male (grey M - Kelly) *pb/pb*.

17 Pb (grey E, litter mate – not actual male) *Pb/pb*.

24 non-Pb (grey) *pb/pb*.

25 non-Pb (McWhite) *pb/pb*.

28 nonPb (blond Ginga) *pb/pb*.

29 Pb (blond Ginga hete) *Pb/pb*.

## Methods

All study fish were raised in 5.75, 8.75 and 10-gallon all-glass aquaria dependent upon age. They received 16 hours of light and 8 hours of darkness per day. Temperatures ranged from 78°F to 82°F. Fish were fed a blend of commercially available vegetable and algae based flake foods and Ziegler Finfish Starter (50/50 mix ratio) twice daily, and newly hatched live Artemia nauplii twice daily. A high volume feeding schedule was maintained in an attempt to produce two positive results: 1. Reduce the time to onset of initial sexual maturity and coloration, thus reduce time between breedings. 2. Increase mature size for ease of phenotypic evaluation and related microscopic study.

## Results

### I. Description and Characteristics: cellular expression of Pb vs. non-Pb

Spectral color is produced by single wavelengths of ambient sunlight. The Visible Wave Length Band (Visual Color) includes: red (*620-670 nm* Bright Red / *670-750 nm* Dark Red), orange and yellow (*570-620 nm*), green (*500-570 nm*), blue (*430-500 nm*), indigo (often omitted in modern times) and violet (*400-430 nm*). Red light, with the longest wavelength and the least amount of energy, allows natural light penetration at less depth. Blue / violet light (*near-UV*), has the shortest wavelength and the most amount of energy, and allows natural light penetration to greater depth. Violet is a true wavelength color, while Purple is a composite effect produced by combining blue and red wavelength colors.

In general, while there are microscopic differences, our findings of visual distinctions between Pb and non-Pb are often more consistent, as opposed to microscopic distinctions. Much of this is likely the result of variability in both zygosity and ornament composition between individuals, both among and between populations and strains. Microscopically, structural differentiation between xantho-erythrophores appears minimal, with differences in population levels and collection or clustering of xanthophores. Heterozygous Pb exhibits partial reduction in collected xanthophores, and homozygous Pb exhibits near complete removal of collected and clustered xanthophores. Though, it is noted yellow color cell populations consisting of isolated “wild-type” single cell xanthophores remain intact.

Dendritic melanophores are present in both Pb and non-Pb at various locations in the body. Observation of Pb in heterozygous and homozygous condition, for mature individuals, reveals that ectopic melanophore dendrites are often extremely extended **(Fig 4)**. This occurs either as the result of direct modification by Pb, or indirectly through interactions as a result of xanthophore reductions or removal (Kottler 2013, 2014, 2015). Overall dendritic melanophore structure is of a much “finer” appearance as compared to non-Pb. Modified melanophores are more often linked together in “chain-like” strings **(Fig 4)**, as compared to non-Pb, both within and outside of areas defined as reticulation along scale edges. While this study did not directly seek to identify an increase in melanophore populations, it was assumed higher numbers of melanophore structures would be present in Pb. While this may be the case, “darker” appearance in Pb vs. non-Pb appears to largely be the result of modification of existing melanophore structures (corolla and punctate) into extended dendrites. Thus, the number of melanophore structures does not appear to drastically increase, in any given individual, only the size of the structures themselves.

**Fig 4.**
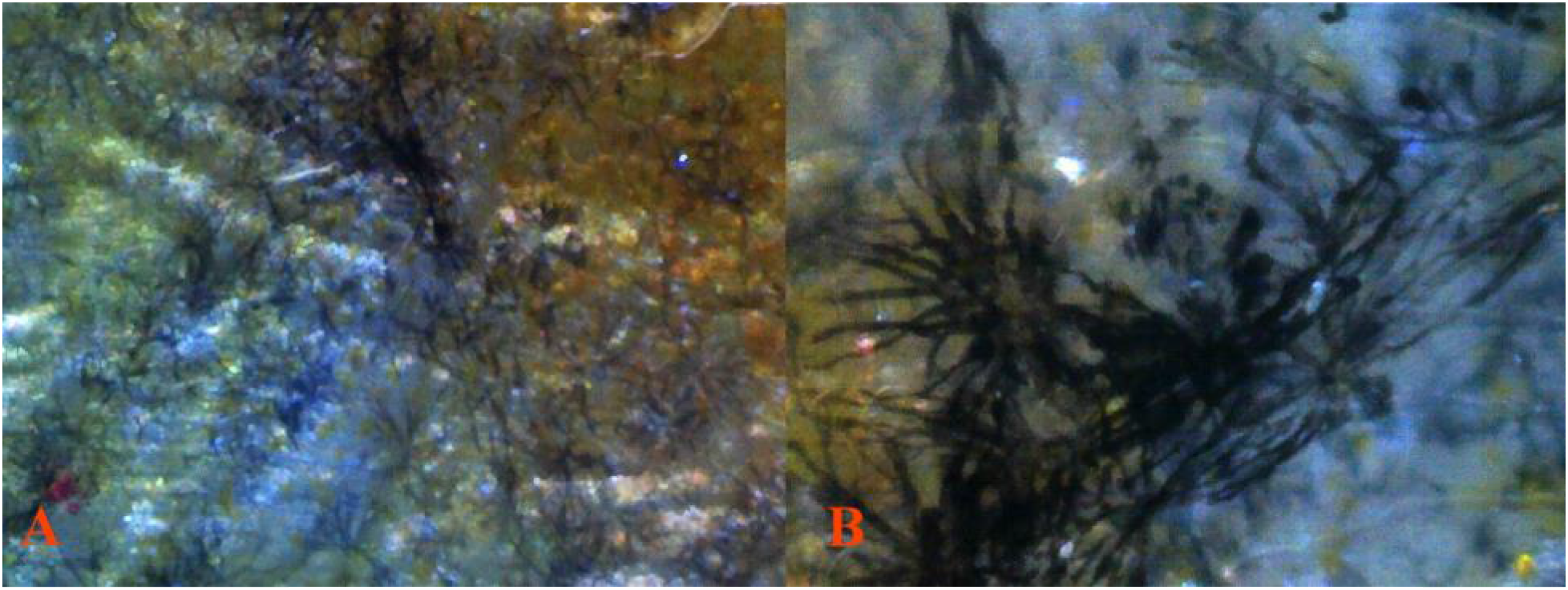
**(A)** 8 Pb 40X 11 under reflected light without transmitted light *(dissected)*. **(B)** 8 Pb 100X 12 under reflected light without transmitted light *(dissected)*. Pb modified dendritic melanophore strings and violet-blue iridophore chromatophore units along scale edging producing reticulated pattern. The extreme “length” of dendrites is a result of Pb.

The motile nature of melanosomes in ectopic melanophores may allow for changes in reflective qualities or hue of individuals. In conjunction with zygosity dependent removal of xantho-erythrophores, this may satisfy both female sexual selective preferences for conspicuous pattern of “bright orange” under specific lighting, and maintain crypsis in others (Endler 1978). While frequent evidence of dendritic and/or motile yellow color pigment (xanthophore) structures was detected in this study, none was found for dendritic and/or motile iridophores, outside of violet and blue [*hereafter violet-blue*] iridophore clustering associated with ectopic dendritic melanophores. Violet-blue iridophores are more visible in Pb vs. non-Pb, with variability between populations and strains, based on overall genotype. Specific to varying genotypes of individuals, there appears to be an actual increase in the ratio of violet to blue iridophores, as would be expected in an “all purple phenotype”. Whether there is an actual increase in iridophore population numbers, or simply increase visibility, due to reductions or removal of xanthophores and/or altered melanophores was not addressed in this study. Nor was the issue of the modification of the angles at which crystalline platelets reside beneath iridophore layers and basal level melanophores.

Prior study of Guppies’ scales, population and strain dependent, show they possess a compliment of chromatophores that include melanophores and xantho-erythrophores (Phang 1985, Ueshima 1998). Their presence and interactions are a component of background body coloration. While iridophores were not specifically mentioned in either study, our work indicates not only their presence in scale epidermis **(Fig 5-6)**, but may indicate increased population numbers in conjunctions with Pb. High numbers of both iridophores and crystalline platelets (Khoo 2010; Bias, *unpublished microscopic observations*) were found on field edges, between circuli and annulus rings of the scales **(Fig 5A** and **Fig 5B)**. Reduced, as a result of Pb modification was the expected minimal number of xanthophores that had been found in non-Pb samples. Erythrophores were not reliably detected in scales of near “wild-type” or feral populations in their study (Phang 1985). These cell types combine to produce not only background body coloration, but also increased reflectivity.

**Fig 5.**
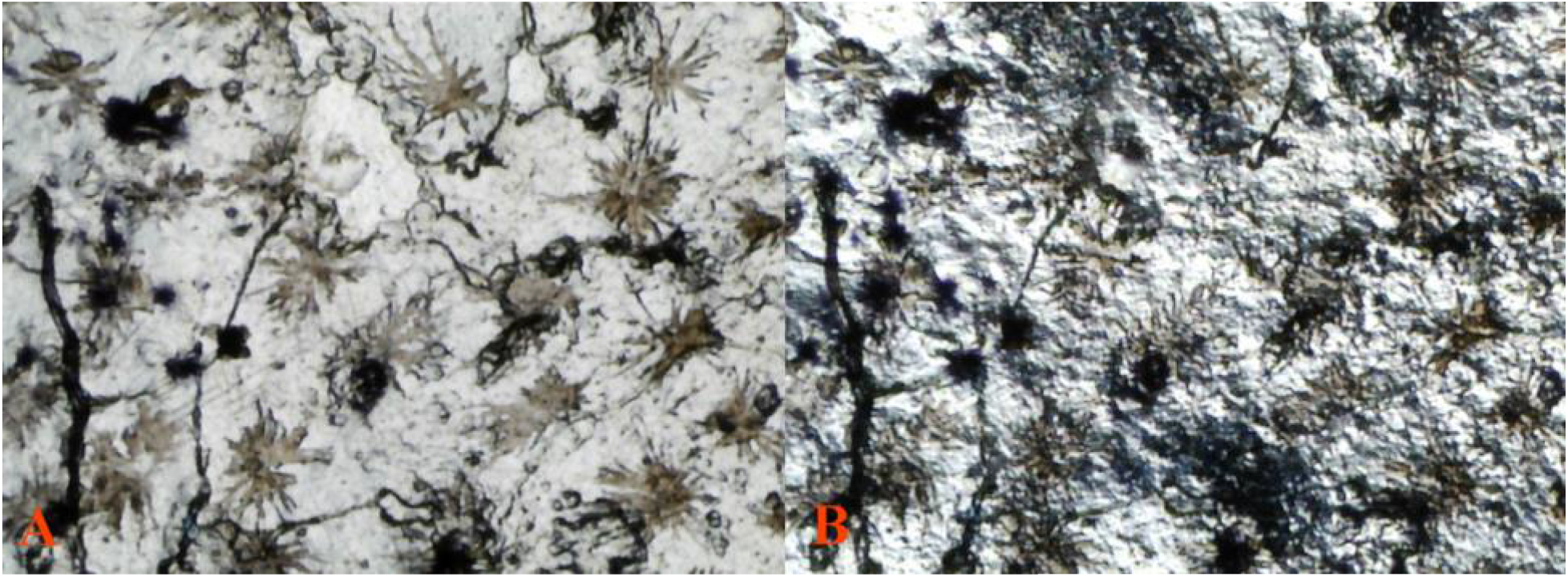
Slide 10 Pb 40X 6 under two light sources. **(A)** 10 Pb 40X 6 scale under transmitted light without reflected light *(dissected*. Lack of reflected light reveals true structure of dendrites with violet-blue iridophores within the organelle, though not reflecting. Variation in color of dendrites shows placement is at varying levels, and organelles are 3-dimentional in composition. **(B)** The same field, under reflected/transmitted light *(dissected)*. Dendritic melanophore-iridophore chromatophore units are visible in the larger structures, violet-blue iridophores laying at lower levels (non-reflecting), and crystalline platelets are all visible.

**Fig 6.**
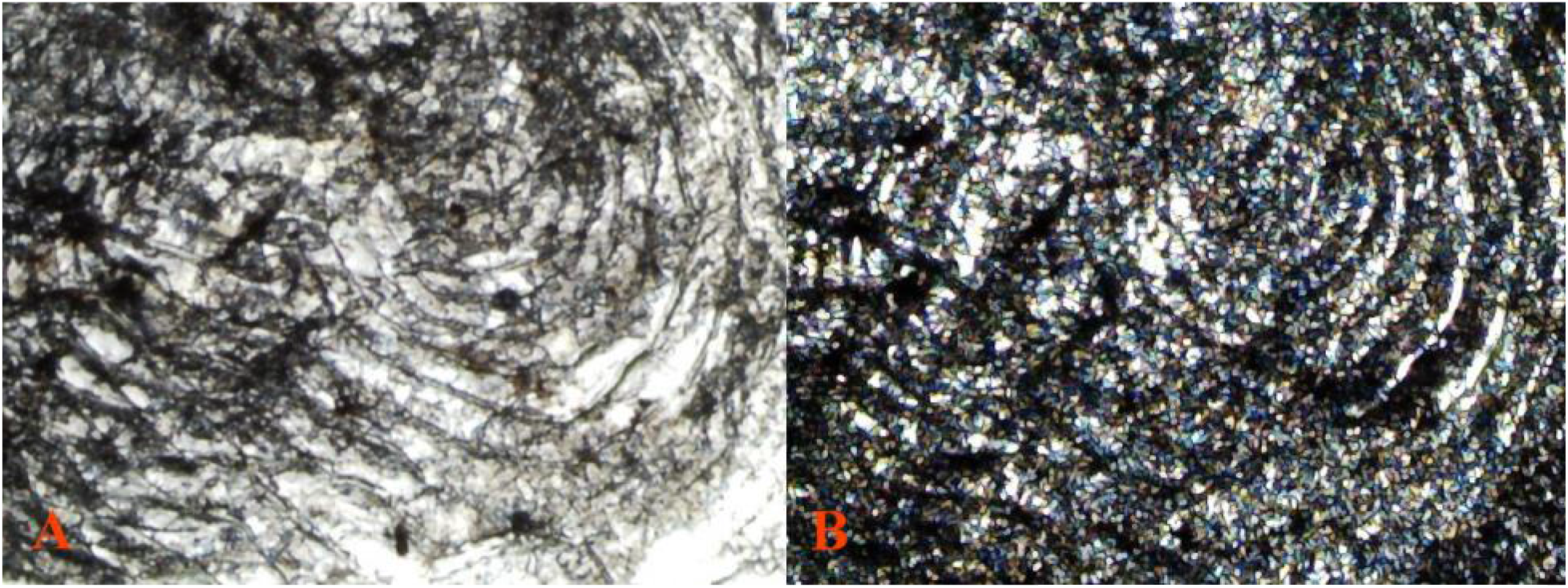
Slide 10 Pb 40X 9 under two light sources. **(A)** 10 Pb 40X 9 scale under transmitted light without reflected light *(dissected)*. Lack of reflected light reveals non-reflective violet-blue iridophores both within the dendrites and scale rings. **(B)** The same field, under reflected/transmitted light *(dissected)*. Presence of violet-blue iridophores, crystalline platelets and minimal color pigments is detected. Focal shifts failed to alter “yellow cell” coloration to either blue or violet.

### II. Cellular Comparison; melanophore modifications in Pb *Pb/pb* vs. non-Pb *pb/pb* under reflected and transmitted light

Dendritic melanophores in heterozygous and homozygous Pb condition, in mature individuals, reveal that dendritic arm structure is extremely extended and finer in appearance **(Fig 7-8)**. Dendrites are linked together in “chain-like” strings intermingled with violet-blue iridophores in chromatophores units. Within the rear peduncle spot and surrounding edges a noticeable absence of corolla and punctate melanophores was often evident. This absence was abated in other regions **(Fig 9-10)** of the body or specific to individuals. The angle of incident lighting and spectral capabilities can alter visual perception, so too can the direction of light. Examples are here presented under both reflected and transmitted lighting, to reveal chromatophore visibility and expression for each.

#### Pb male posterior peduncle from the caudal base to just below the dorsal base

**Fig 7.**
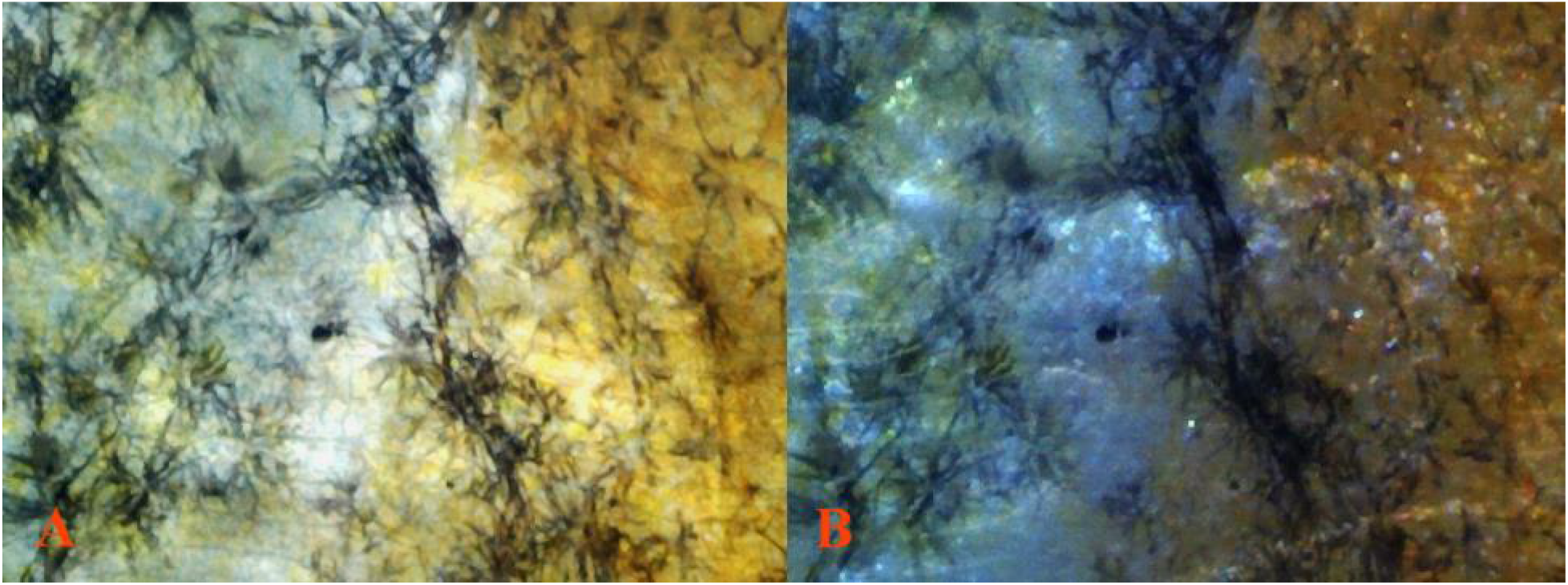
**(A)** 8 Pb 40X 13 *Pb/pb (dissected)* transmitted light. **(B)** The same field, reflected light. Extreme dendritic melanophore modification in reticulated pattern do to heterozygous Pb.

**Fig 8.**
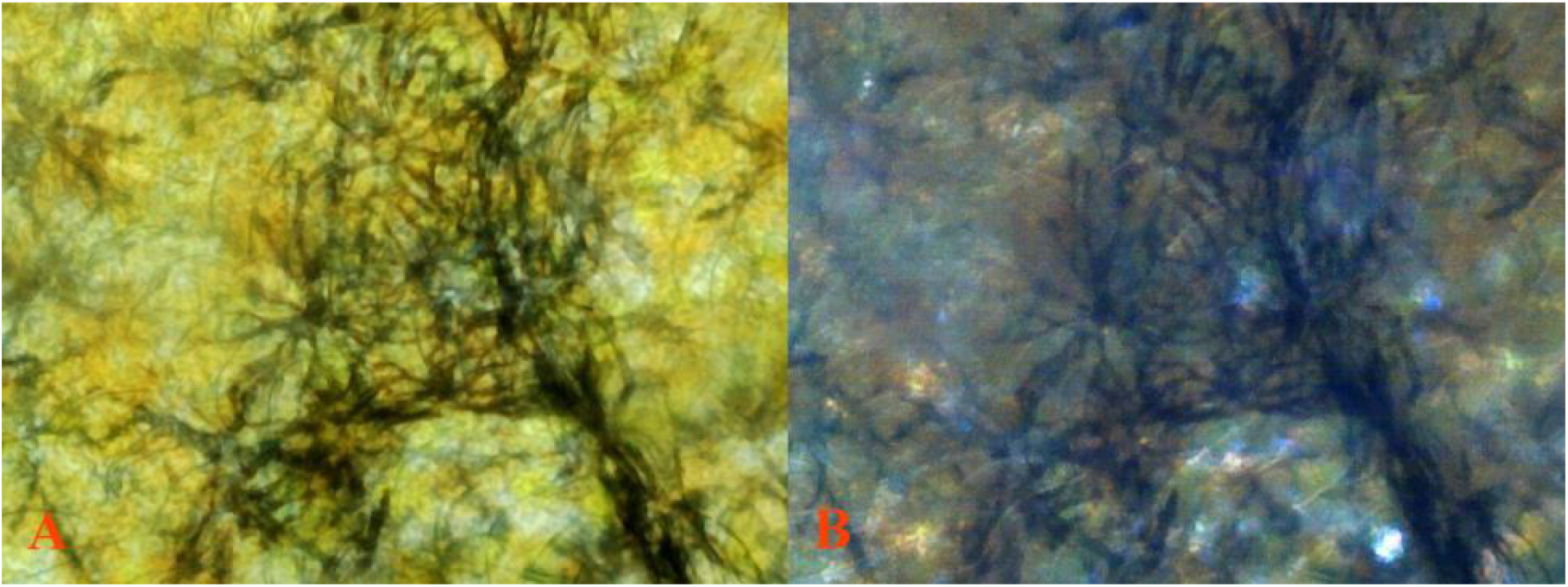
**(A)** 8 Pb 100X 2 *Pb/pb (dissected)* transmitted light. **(B)** The same field, reflected light. Extreme dendritic melanophore modification in reticulated pattern by heterozygous Pb.

**Fig 9.**
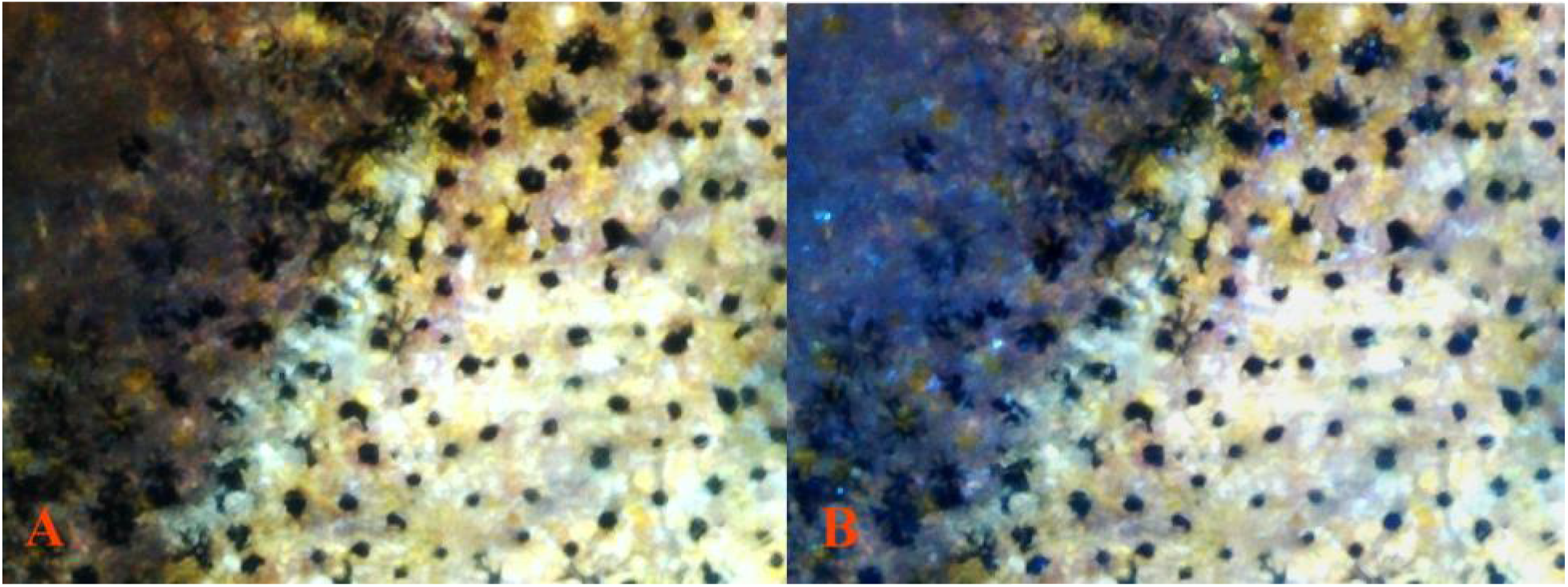
**(A)** 2 non-Pb 40X 11 *pb/pb (dissected)* transmitted light. **(B)** The same field, *(dissected)* reflected/transmitted light.

**Fig 10.**
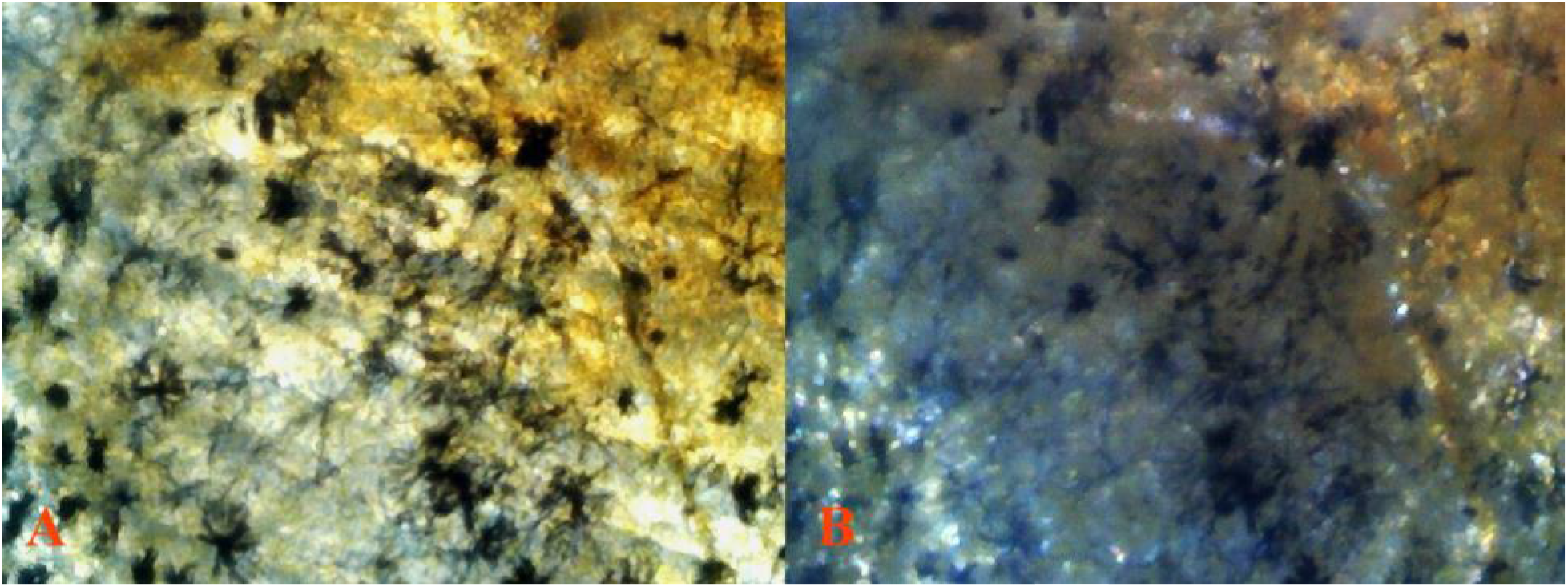
**(A)** 7 non-Pb 40X 15 *pb/pb (dissected)* transmitted light. **(B)** The same field, *(dissected)* reflected light. Reduced dendritic melaophore extensions in non-Pb.

#### Non-Pb male posterior peduncle from the caudal base to just below the dorsal base

### III. Cellular Comparison: early coloration in Pb *Pb/pb* vs. non-Pb *pb/pb* male Guppies (*Poecilia reticulata*)

The following examples of early coloration in non-Pb and Pb show male expression of violet-blue iridophores macroscopically **(Fig 11)** and in 100x and 400x **(Fig 12-15)**. Visual distinction is easily made between the two iridophore types. Changes in magnification, progressive focal shift, adjustment in angle of incident lighting or direction of light (reflected or transmitted) consistently failed to remove this visible distinction between violet and blue. Thus, this demonstrates two distinct iridophore populations in the blue-violet spectrum.

**Fig 11.**
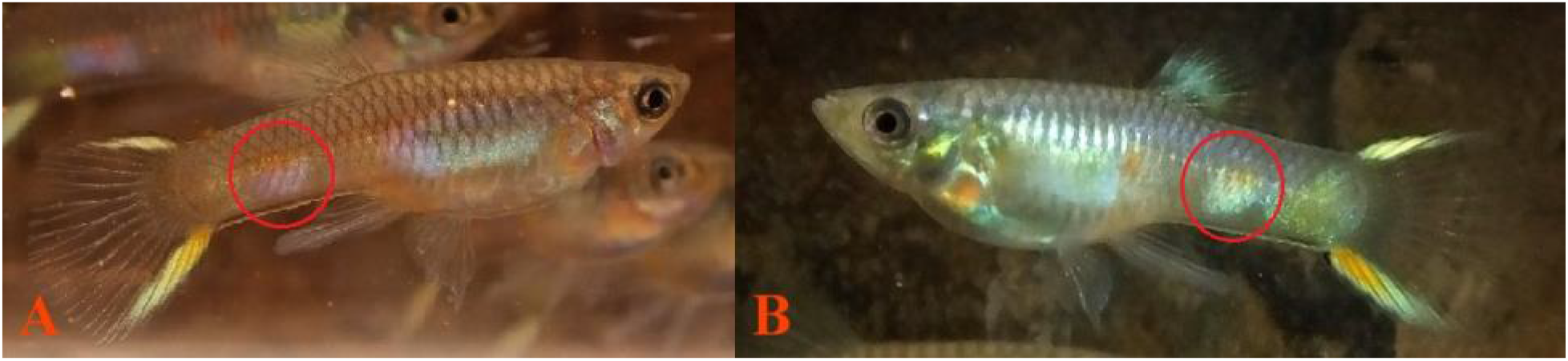
**(A)** 6 Pb male (grey) *Pb/pb*, **(B)** 3 non-Pb male (grey) *pb/pb*. All slide images taken from posterior orange spot (red circle).

**Fig 12.**
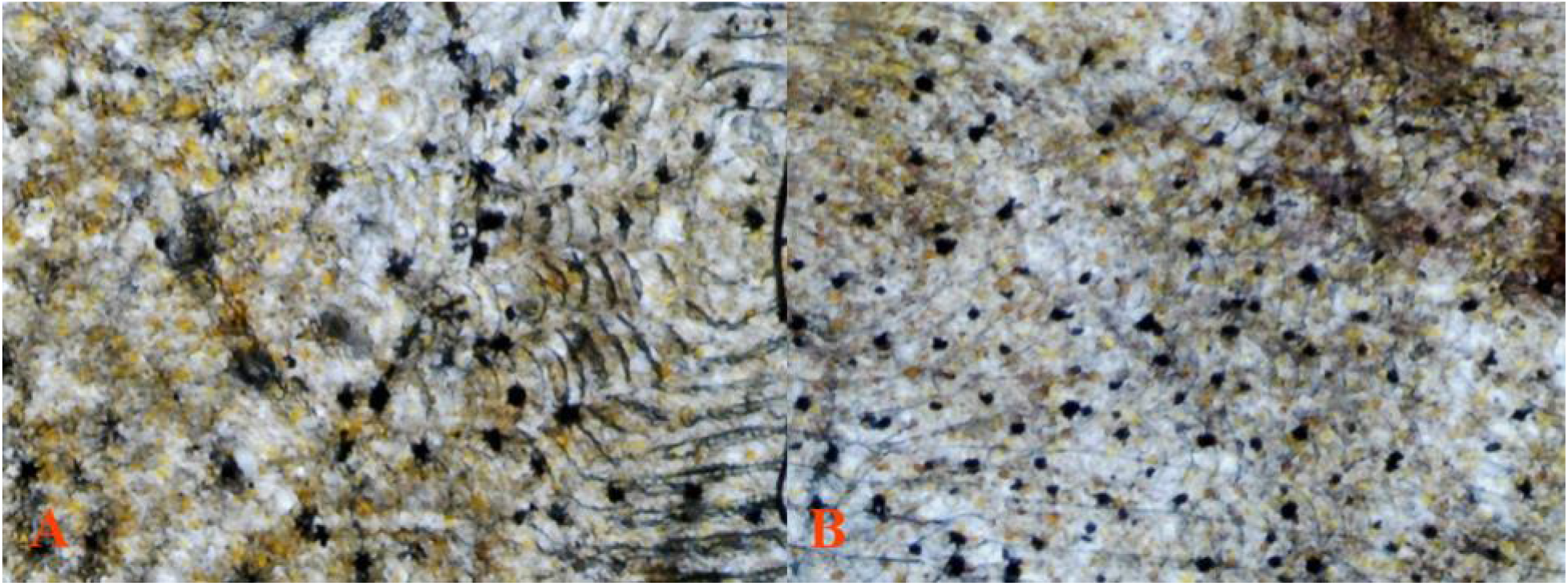
**(A)** 6 Pb 40X 14 *Pb/pb (dissected)* transmitted light. Increased dendritic melanophore extensions in Pb at similar age. **(B)** 3 non-Pb 40X 12 *pb/pb (dissected)* transmitted light.

**Fig 13.**
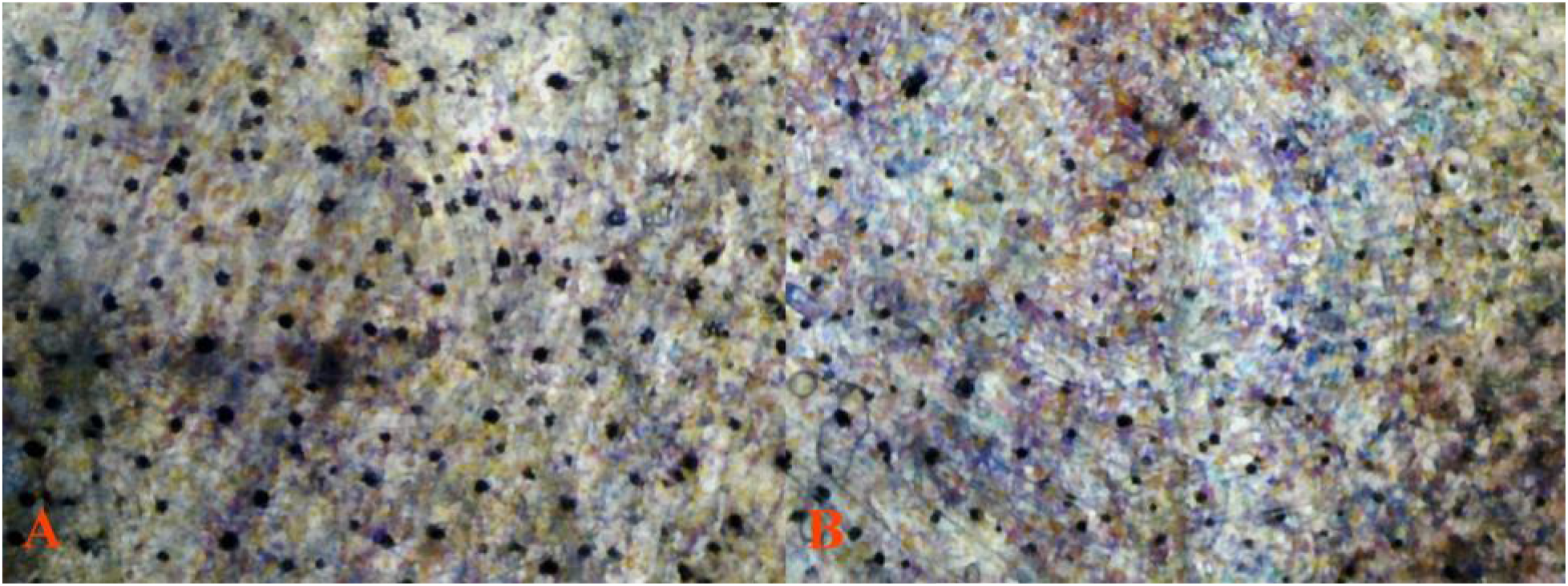
**(A)** 6 Pb 40X 8 *Pb/pb (dissected)* transmitted light. Increased dendritic 267 melanophore extensions in Pb at similar age. **(B)** 3 non-Pb 40X 4 *pb/pb (dissected)* 268 transmitted light. More evenly distributed blend of violet and blue irdophores in non-Pb as 269 compared to Pb.

**Fig 14.**
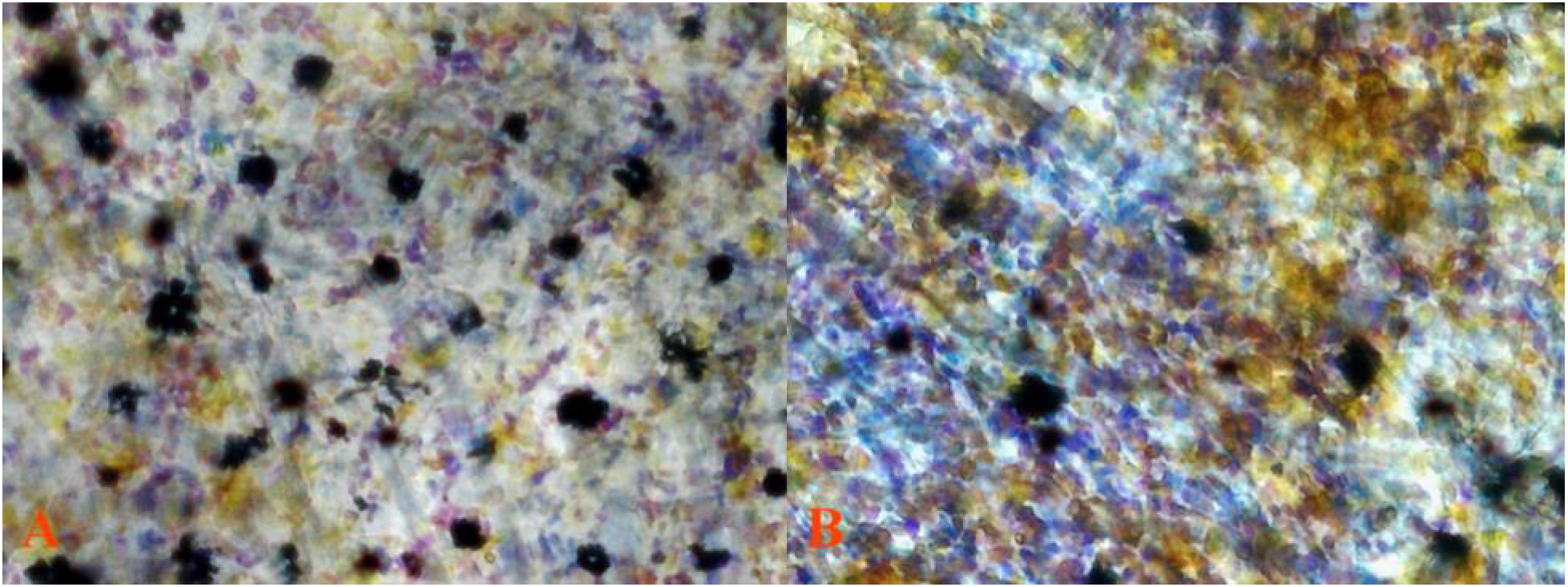
(**A**) 6 Pb 100X 12 *Pb/pb (dissected)* transmitted light. More violet iridophores than 273 blue irdophores. (**B**) 3 non-Pb 100X 14 *pb/pb (dissected)* transmitted light. More evenly 274 distributed blend of violet and blue irdophores in non-Pb as compared to Pb.

**Fig 15.**
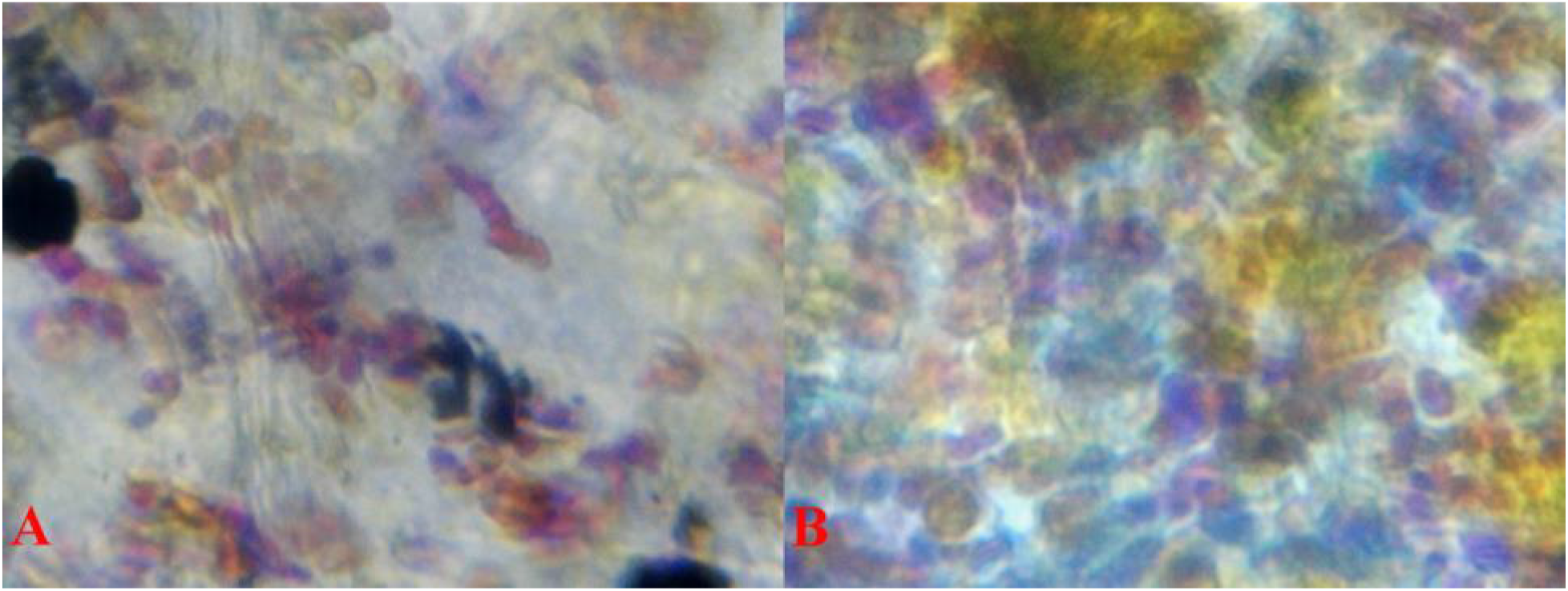
**(A)** 6 Pb 100X 3 *Pb/pb (dissected)* transmitted light. More violet iridophores than blue irdophores. **(B)** 3 non-Pb 100X 6 *pb/pb (dissected)* transmitted light. More evenly distributed blend of violet and blue irdophores in non-Pb as compared to Pb.

Two distinct observations are offered based on early coloration. First, violet and blue iridophores appear “randomly collected among themselves” in similar fashion **(Fig 13-15)**, as opposed to later mature coloration in which violet and blue iridophores are arranged together in “joined alternating color” groupings in dissimilar fashion. This shows that coloration is nearly complete, while migration to their final location is not. Second, melanophore shape is predominately corolla or punctate in early coloration **(Fig 12-14)**, as opposed to mature coloration in which dendrites dominate. This indicates that members of the melanophore population are in place, while their final shape is not established. Side by side presentation of similar locations in Pb and non-Pb are presented.

### IV. Cellular Comparison: late coloration in Pb *Pb/pb* vs. non-Pb *pb/pb* male Guppies (*Poecilia reticulata*)

Macroscopically **(Fig 16)** and microscopically **(Fig 17-21)** visible in heterozygous Pb are partial reductions in collected xanthophores, and in homozygous Pb near complete removal of collected and clustered xanthophores. Yellow color cell populations consisting of isolated “wild-type” single cell xanthophores remain intact.

**Fig 16.**
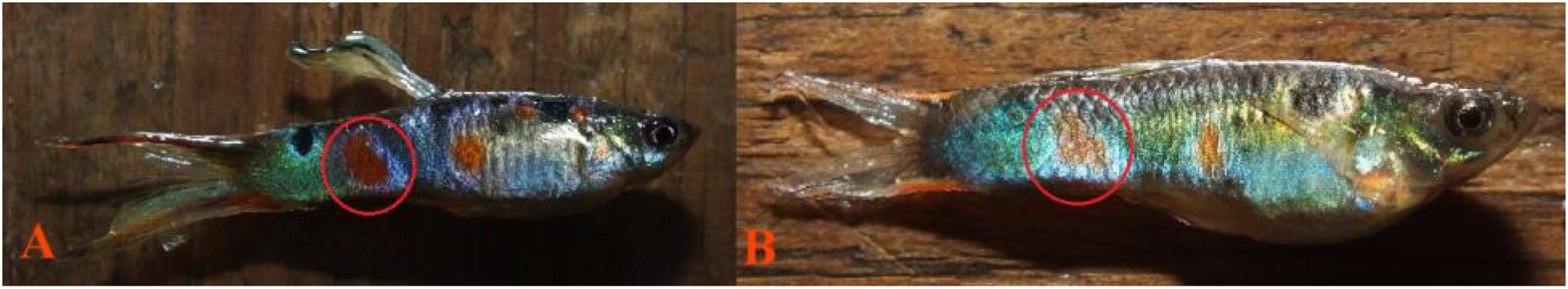
**(A)** Heterozygous 11 Pb male *Pb/pb*, **(B)** 6 non-Pb male *pb/pb*. All slide images taken from posterior orange spot (red circle).

**Fig 17.**
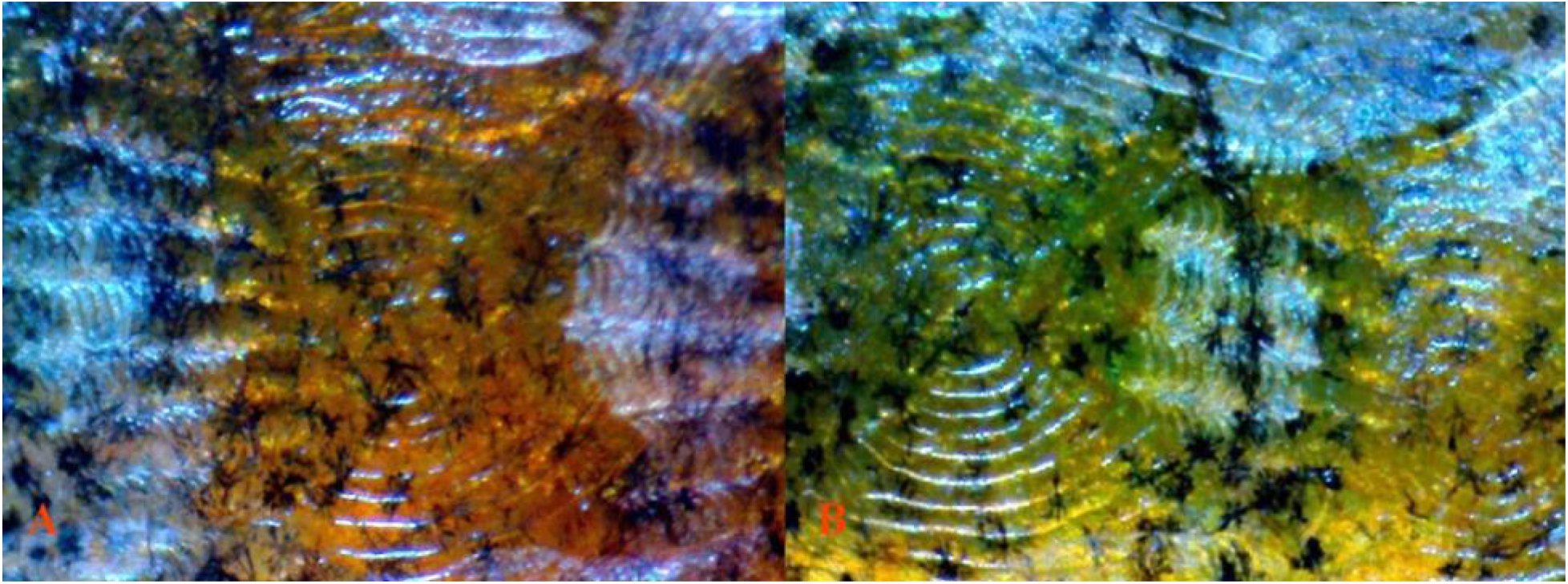
**(A)** 13 Pb 40X 4 *Pb/Pb (dissected)* reflected light. Expected higher visiblity of 321 erythrophores with reduction of xanthophores by Pb modification. **(B)** 14 non-Pb 40X 3 322 *pb/pb (dissected)* reflected light. Expected evenly distributed xantho-erythrophores in non-323 Pb.

**Fig 18.**
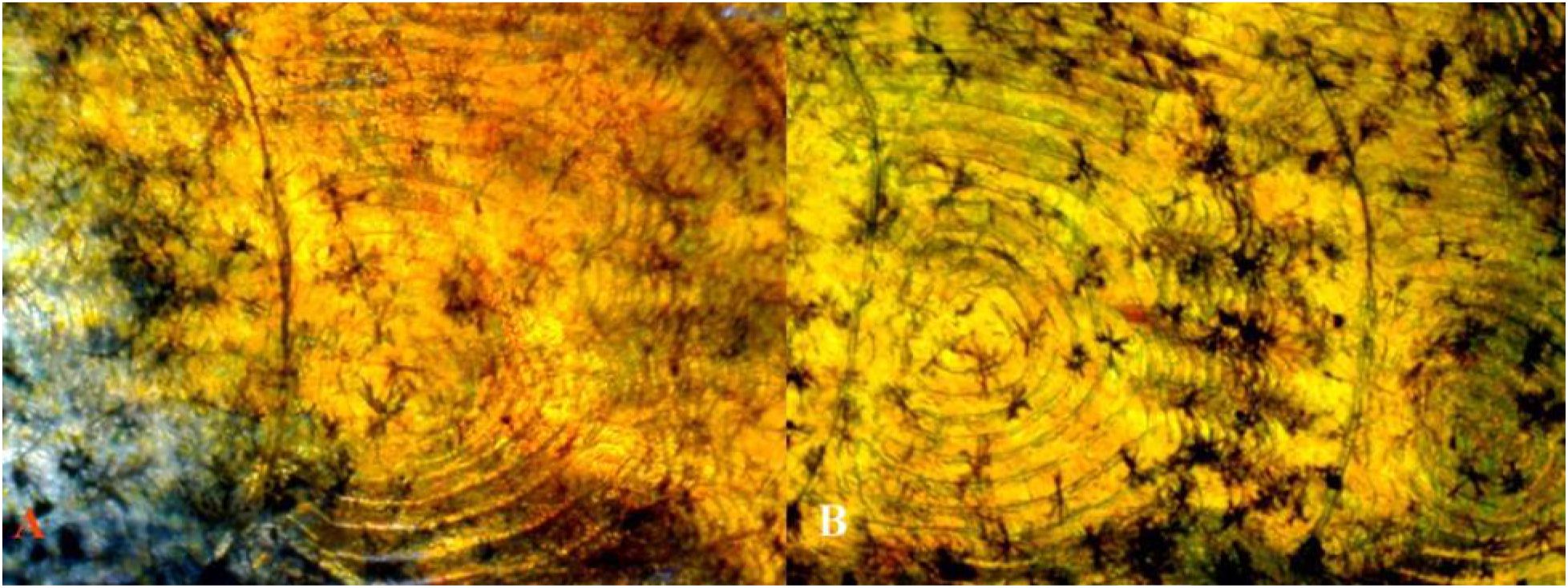
**(A)** 13 Pb 40X 4 *Pb/Pb (dissected)* reflected/transmitted light. Expected higher visiblity of erythrophores with reduction of xanthophores by Pb modification. **(B)** 14 non-Pb 40X 3 *pb/pb (dissected)* transmitted light. Expected evenly distributed xantho-erythrophores in non-Pb.

**Fig 19.**
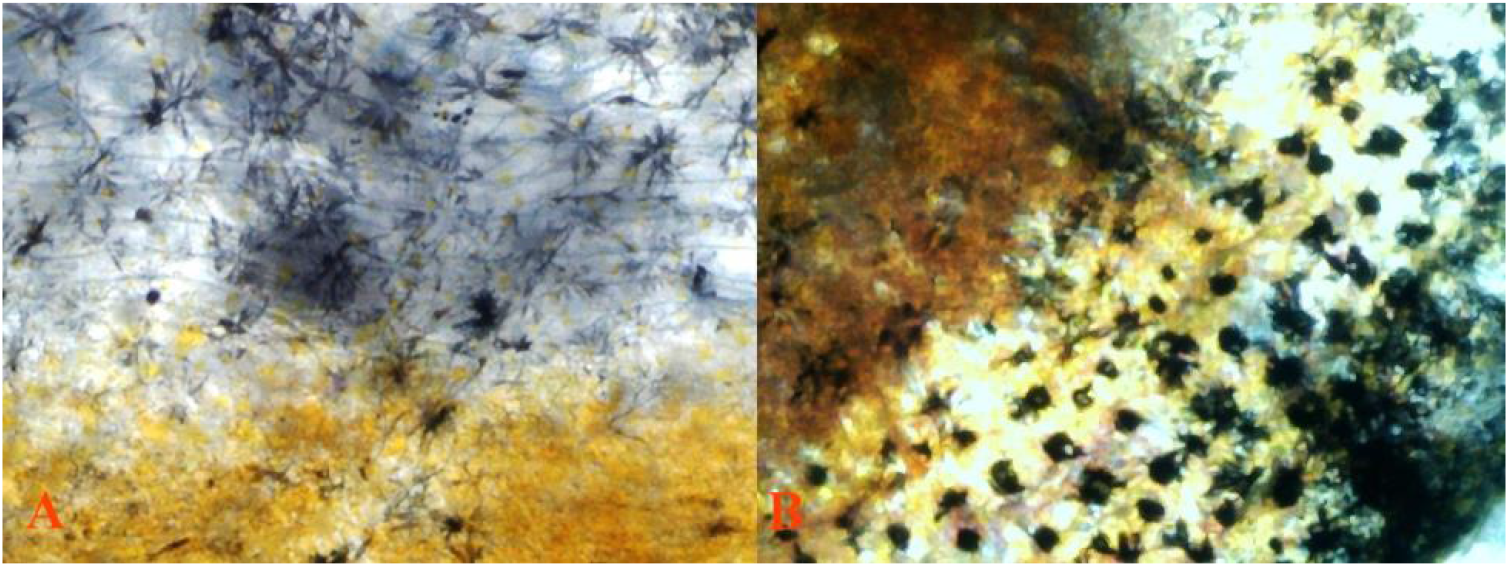
**(A)** 2 Pb 40X 1 *Pb/pb (dissected)* transmitted light. Extreme dendritic melanophore modification by Pb. **(B)** 2 non-Pb 40X 4 *pb/pb (dissected)* transmitted light. Minimal dendritic melanophore modification by non-Pb.

**Fig 20.**
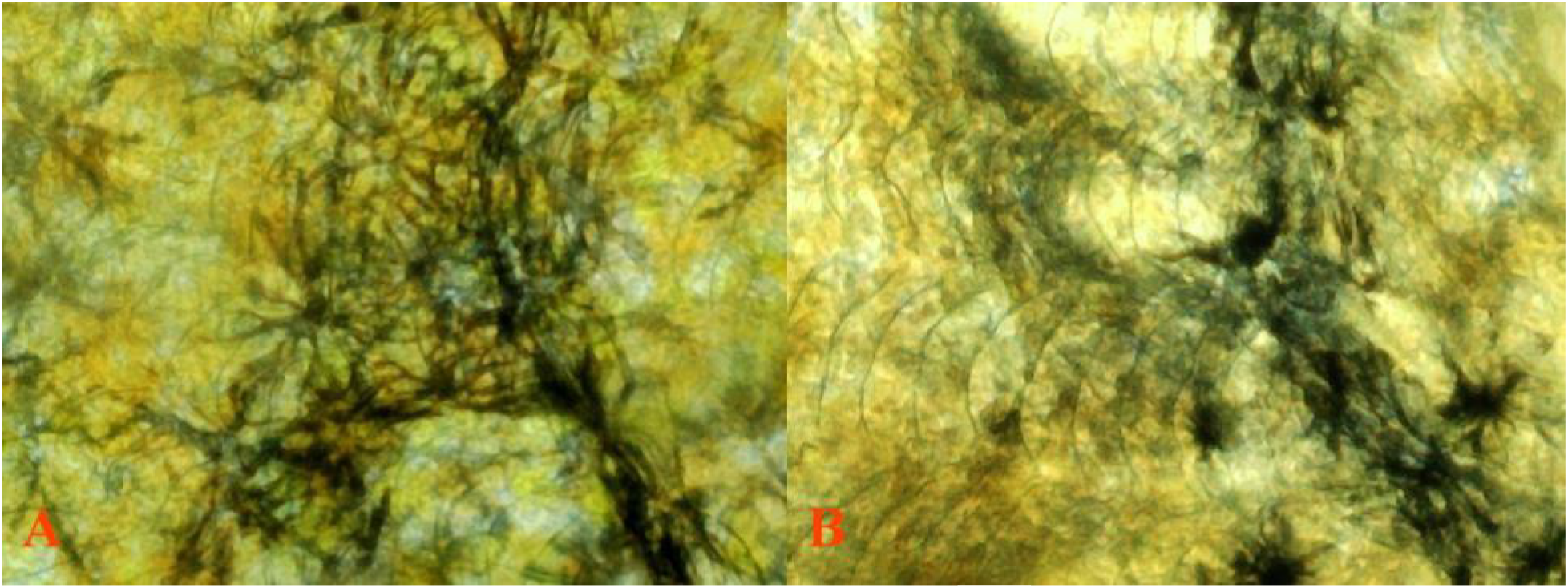
**(A)** 8 Pb 100X 2 *Pb/pb (dissected)* transmitted light. Extreme dendritic 338 melanophore modification by Pb. **(B)** 7 non-Pb 100X 17 *pb/pb (dissected)* transmitted 339 light. Minimal dendritic melanophore modification by non-Pb.

**Fig 21.**
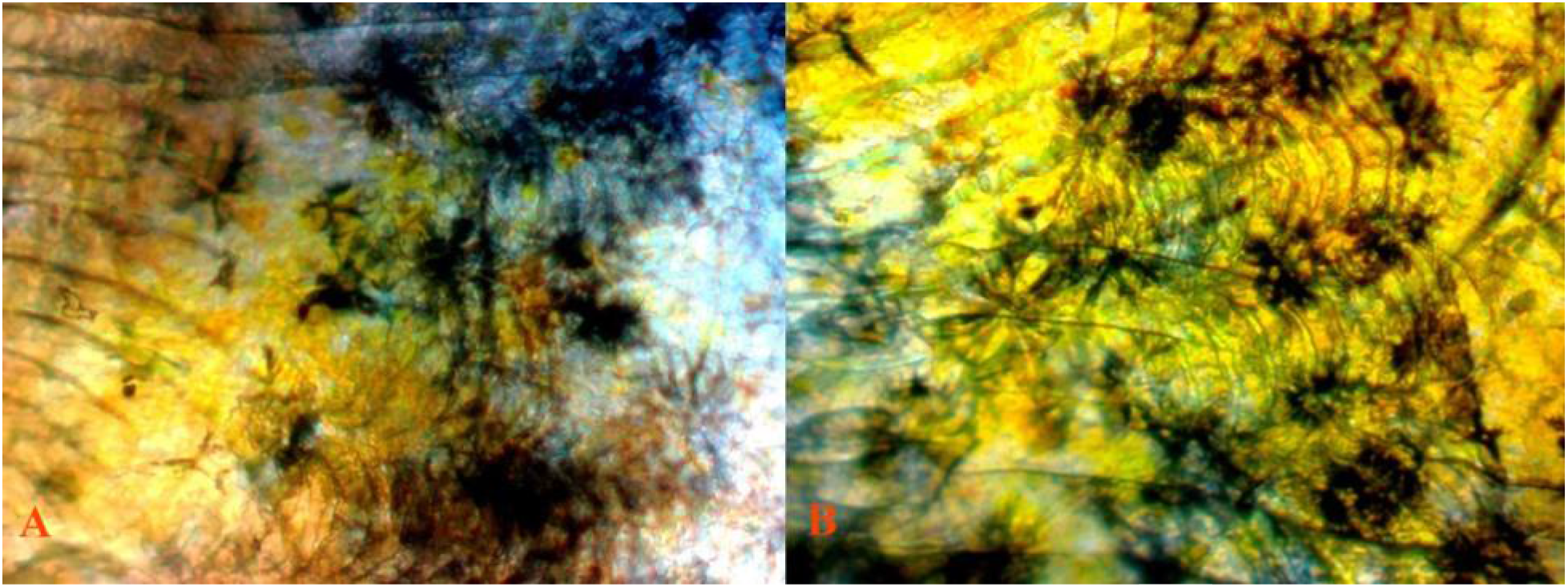
**(A)** 13 Pb 100X 4 *Pb/Pb (dissected)* reflected/transmitted light. Expected higher 343 visiblity of erythrophores with reduction of xanthophores by Pb modification. **(B)** 14 non-344 Pb 100X 3 *pb/pb* reflected/transmitted light. Expected evenly distributed xantho-345 erythrophores in non-Pb.

Dendritic melanophores in heterozygous and homozygous Pb condition, for mature individuals, reveal that dendrite structure is extremely extended and finer in appearance **(Fig 19-20)**. Dendrites are linked together in “chain-like” strings intermingled with violet-blue iridophores in chromatophores units, while corolla melanophores are present in lessor numbers and punctate melanophores nearly absent **(Fig 19)**. Darker appearance, both microscopically and phenotypically, results from modification of existing melanophore structures into extended dendrites.

All major classes of chromatophores were present in the rear peduncle spot and adjoining areas in both Pb and non-Pb. Violet-blue iridophores are more visible in Pb vs. non-Pb, with variability between study specimens. An increase in the ratio of violet to blue iridophores was observed. Collected and clustered xanthophore populations, found in non-Pb members of the contemporary group, were reduced in heterozygous Pb condition and removed in homozygous Pb condition. The retention of isolated xanthophores remained intact in both heterozygous and homozygous Pb condition.

Violet and blue iridophores appear arranged together in “joined alternating color” groupings in dissimilar fashion, as opposed to early mature coloration in which violet and blue iridophores appear “randomly collected among themselves”. This indicates that coloration and migration are complete. Dendritic melanophores dominate shape, as opposed to early coloration in which corolla or punctate melanophores outnumber dendritic melanophores. This indicates that population numbers and shape are in place. Side by side presentation of similar locations in Pb and non-Pb are presented.

### V. Cellular Comparison: late coloration in Asian Blau *Ab/ab Pb/Pb* vs. non-Pb *Ab/ab pb/pb* male Guppies (*Poecilia reticulata*)

Asian Blau (*Ab* – *undescribed*, see Bias 2015) presents a unique opportunity to further confirm the spectral removal of yellow color pigment by Pb, though microscopic study reveals this removal to be far from complete. Autosomal incompletely dominant Ab, as opposed to autosomal recessive European Blau (*r* or *r1*, Dzwillo 1959) in heterozygous and homozygous condition removes red color pigment (**Fig 22A-B**).

**Fig 22.**
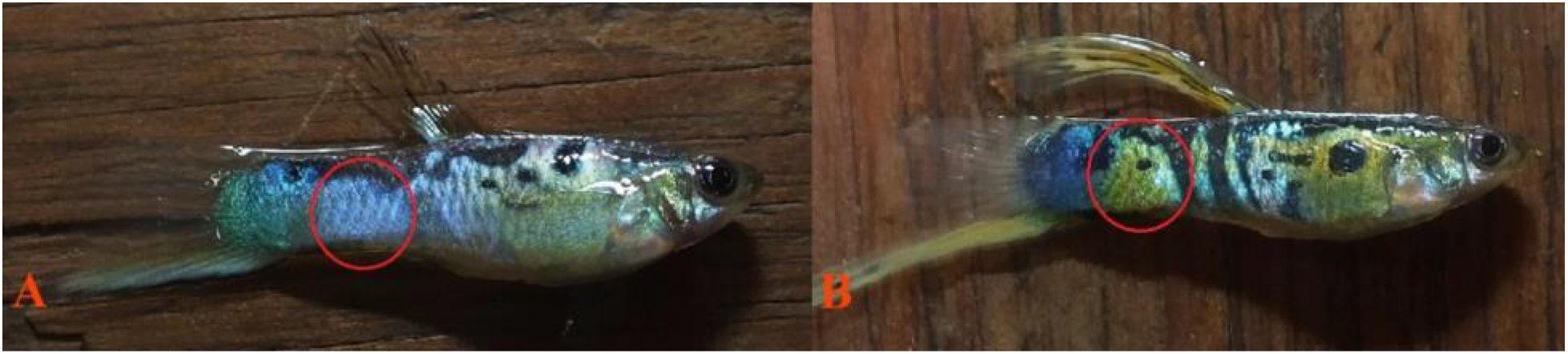
**(A)** 5 Pb male (grey) *Pb/Pb Ab/ab*, **(B)** 4 non-Pb male (grey) *pb/pb Ab/ab*. All 372 slide images taken from posterior modified orange spot (red circle).

Collected yellow color pigment and clustered Metal Gold (*Mg* - *undescribed, see Bias 2015*) xanthophores are little affected by this erythrophore defect **(Fig 22B),** as shown in the *pb/pb Ab/ab* male. The following macroscopic photo clearly reveals near complete removal of densely packed collected yellow cells in the *Pb/Pb Ab/ab* **(Fig 22A)** male, leaving an underlying “circular ring” of violet-blue iridophores intact. As previously noted, Pb in itself has little or no effect on erythrophore populations. Albeit, Pb modification results in increased expression of violet iridophores (**Fig 22A** vs. **22B**).

The macroscopic presence of underlying iridophores, lacking a xantho-erythrophore (yellow-orange) overlay in *Pb/Pb Ab/ab* **(Fig 22A)** and lacking erythrophore (orange) overly in non-Pb *pb/pb Ab/ab* **(Fig 22B)**, allows for the visual distinction between xantho-erythrophore populations.

Though structural differences between xanthophores and erythrophores may be limited to variability in placement of underlying reflective crystalline platelets (Kottler 2014), microscopically the prior results are confirmed in comparison. Dendrites remain extremely extended and linked together in “chain-like” strings intermingled with violet-blue iridophores in chromatophore units. A side by side comparison of similar locations for Pb *(Pb/Pb Ab/ab)* and non-Pb *(pb/pb Ab/ab)* is presented **(Fig 23-24**).

**Fig 23.**
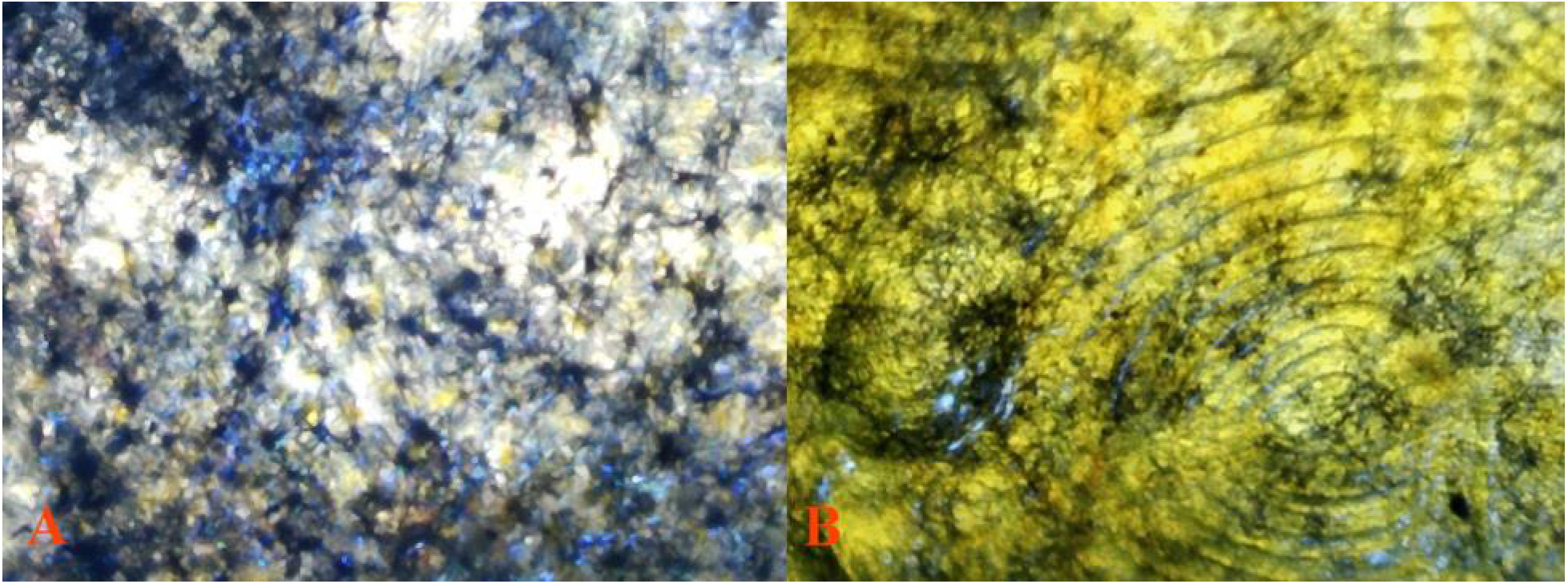
**(A)** 5 Pb 40X 16 *Pb/Pb Ab/ab (dissected)* reflected/transmitted light. Underlying 384 violet-blue iridophore structure is clearly revealed in absence of collected xantho-385 erythrophores. **(B)** 4 non-Pb 40X 8 *pb/pb Ab/ab (dissected)* reflected/transmitted light. Collected xanthophores masking violet-blue iridophores in absence of erythrophores.

**Fig 24.**
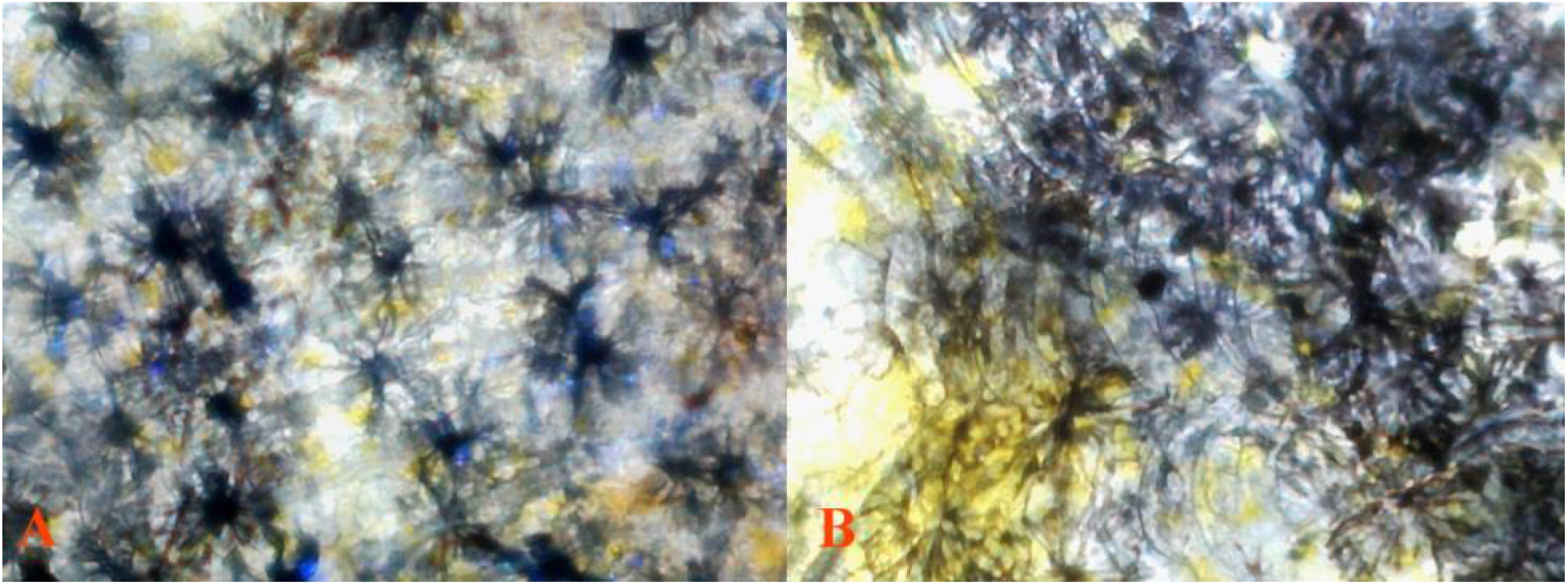
**(A)** 5 Pb 100X 1 *Pb/Pb Ab/ab (dissected)* reflected/transmitted light. Dendritic melanophore modification by homozygous Pb. Composition of dendritic melanophore-iridophore chromatophore units visible. **(B)** 4 non-Pb 100X 9 *pb/pb Ab/ab (dissected)* reflected/transmitted light. Collected xanthophores masking violet-blue iridophores in absence of erythrophores.

### VI. Cellular Comparison: late coloration in Blond *bb, Pb/pb* vs. Blond non-Pb *bb, pb/pb* and interactions with Asian Blau *Ab* male Guppies (*Poecilia reticulata*)

A blond (*bb*) interaction with both Purple Body (*Pb*) and Asian Blau (*Ab*) at the macroscopic level serves two purposes. First, the same general observations are again confirmed macroscopically for removal of collected xanthophores by Pb in body and finnage. Resulting in a modification of orange ornaments to a “pinkish-purple” coloration, with an increase in violet iridophores and overall reflective qualities **(Fig 25A** vs. **25B)**. Second, removal of erythrophores by Ab again confirms nearly complete removal of densely packed collected yellow cells with an increase in violet iridophores by Pb in the *bb, Pb/pb Ab/ab* **(Fig 26A)** male, leaving an underlying “circular ring” of violet-blue iridophores intact. In contrast collected yellow color pigment and clustered Mg xanthophores are little affected by the Ab erythrophore defect **(Fig 26B),** as shown in the *bb, pb/pb Ab/ab* male with no increase in violet iridophores.

**Fig 25.**
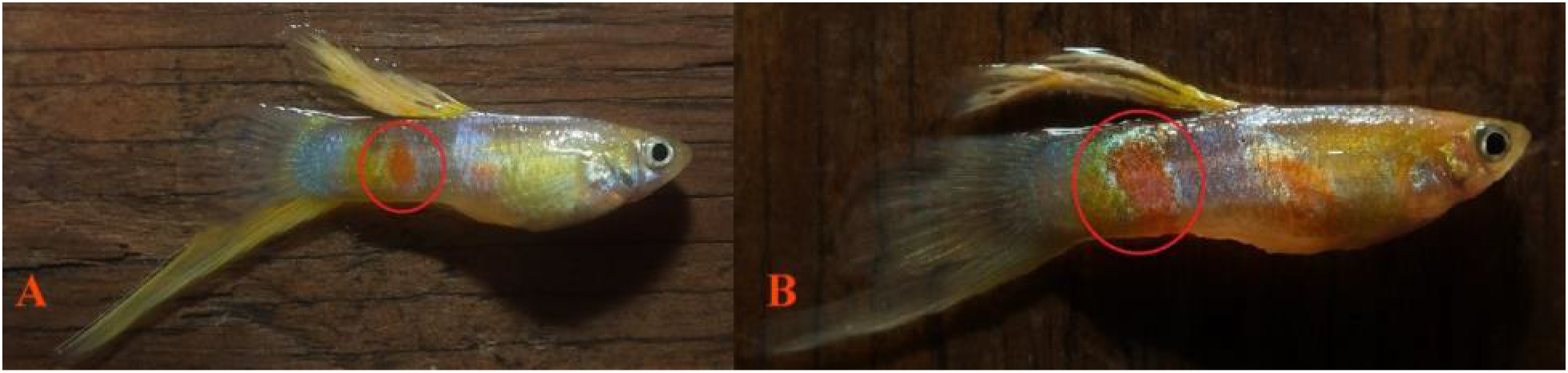
**(A)** 7 Pb male (blond) *bb, Pb/pb*, **(B)** 5 non-Pb male (blond) *bb*, pb/pb. All slide images taken from posterior orange spot (red circle).

**Fig 26.**
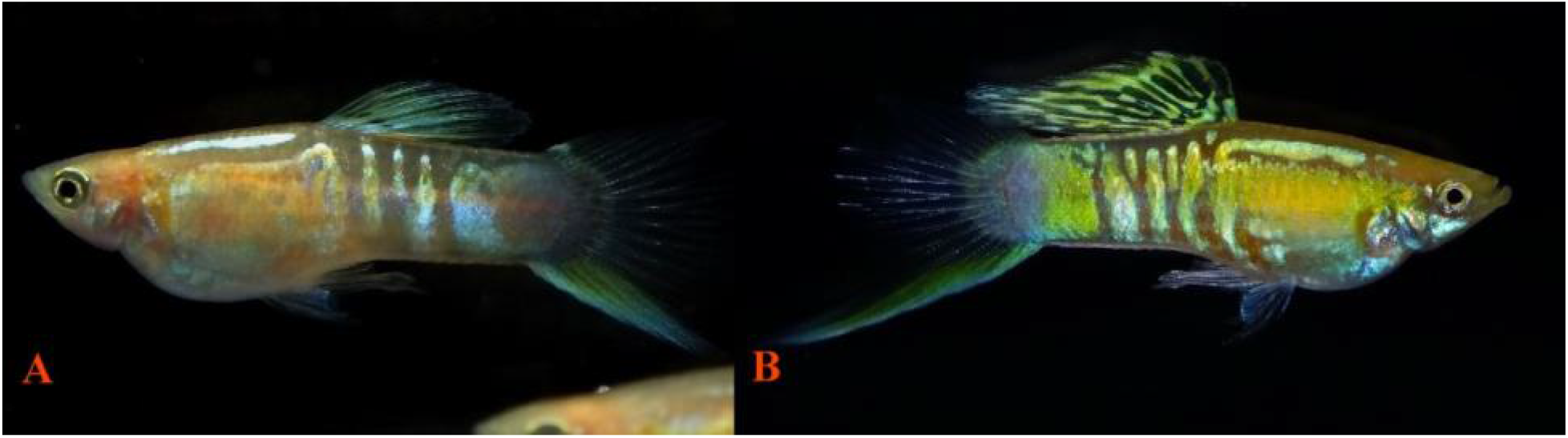
**(A)** Pb male (blond) *bb, Pb/pb Ab/ab*, **(B)** non-Pb male (blond) *bb, pb/pb Ab/ab*.

Modification by Blond results in near normal population levels of melanophores, while they are reduced in size. Blond Pb corolla melanophores are similarly clumped together along scale edging or isolated groups, in the equivalent of grey Pb “chain-like” strings, and overlay violet-blue iridophore chromatophore units. Microscopically, it is noted that the presence of large Pb modified melanophore dendrites is not required for expression of modified “pinkish-purple” ornaments; rather only “brightness” is altered by their reduction. A side by side comparison of similar locations for Pb *bb, Pb/Pb Ab/ab* and non-Pb *bb, pb/pb Ab/ab* is presented **(Fig 27-29)**.

**Fig 27.**
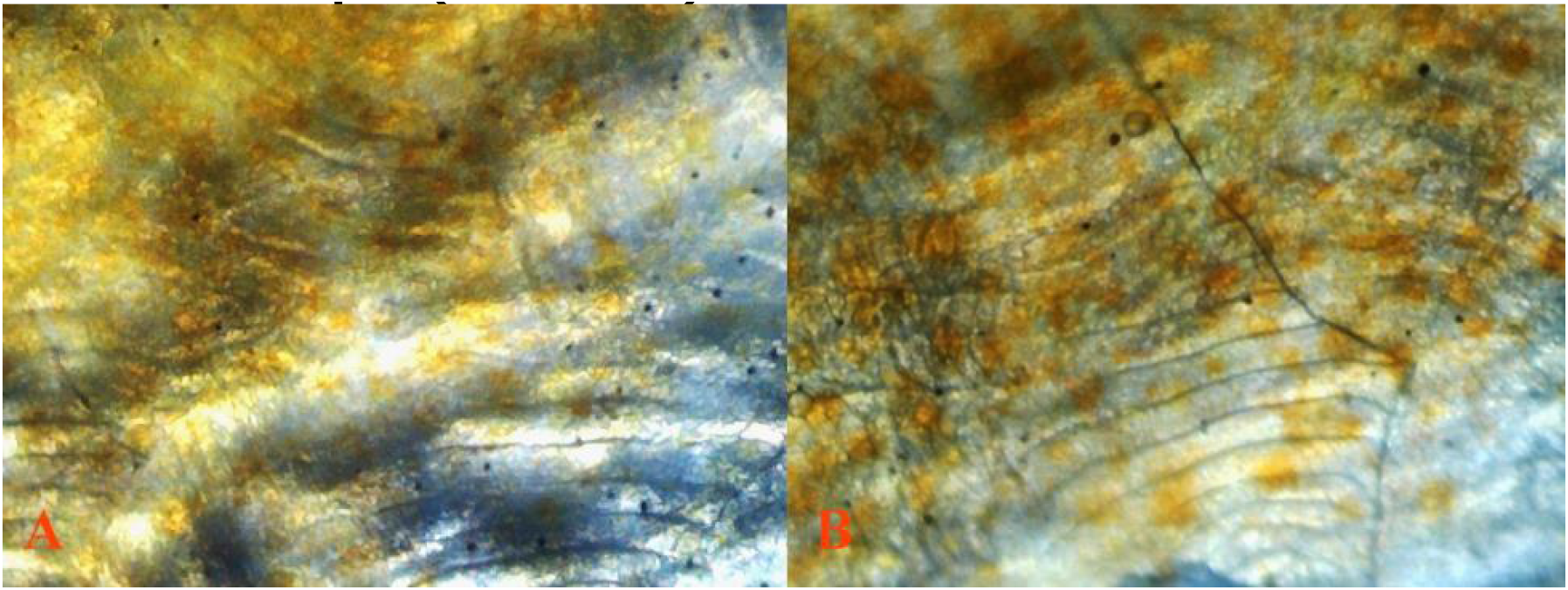
**(A)** 7 Pb 40X 16 *bb, Pb/pb* (*dissected*) transmitted light. Minimally dendritic melanophores are present. **(B)** 5 non-Pb 40X 3 *bb, pb/pb* (*dissected*) transmitted light.

**Fig 28.**
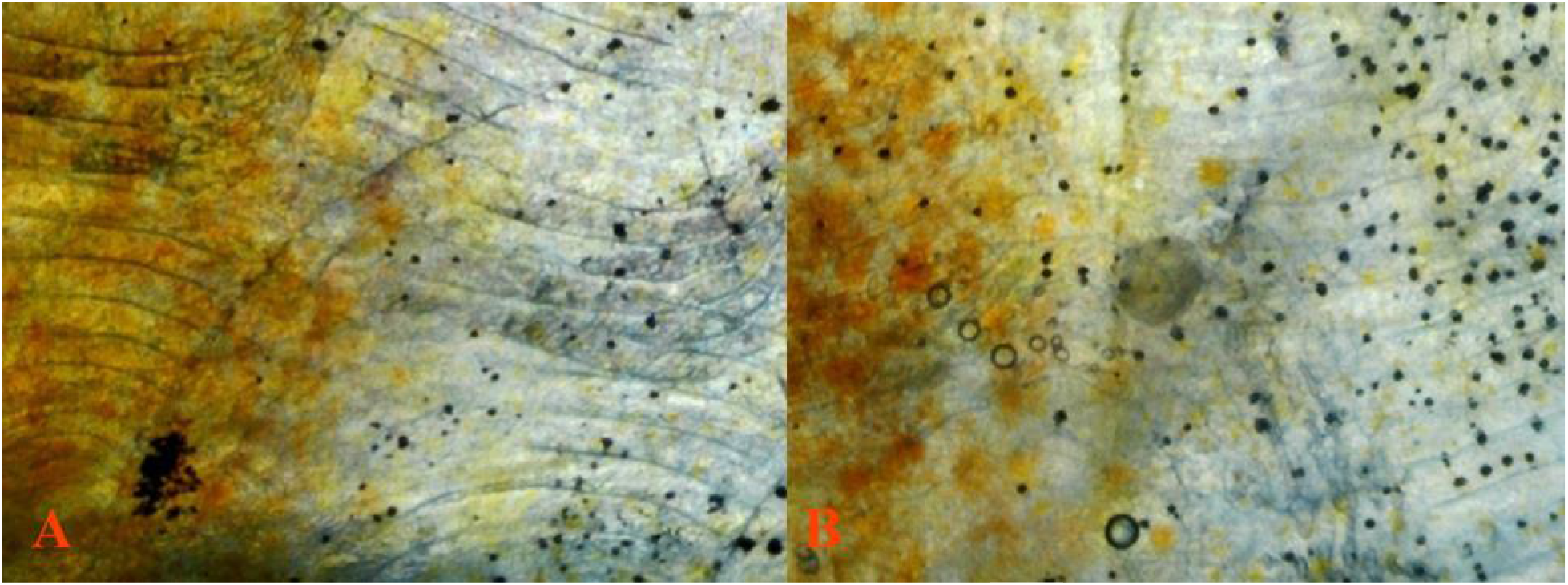
**(A)** 7 Pb 40X 6 *bb, Pb/pb* (*dissected*) transmitted light. Expected higher concentration of erythrophores present with reduced xanthophores from Pb modification. Erythrophores generally more visible from transmitted light. **(B)** 5 non-Pb 40X 4 *bb, pb/pb (dissected)* transmitted light. Expected evenly distributed xantho-erythrophores present in non-Pb. Erythrophores generally more visible from transmitted light.

**Fig 29.**
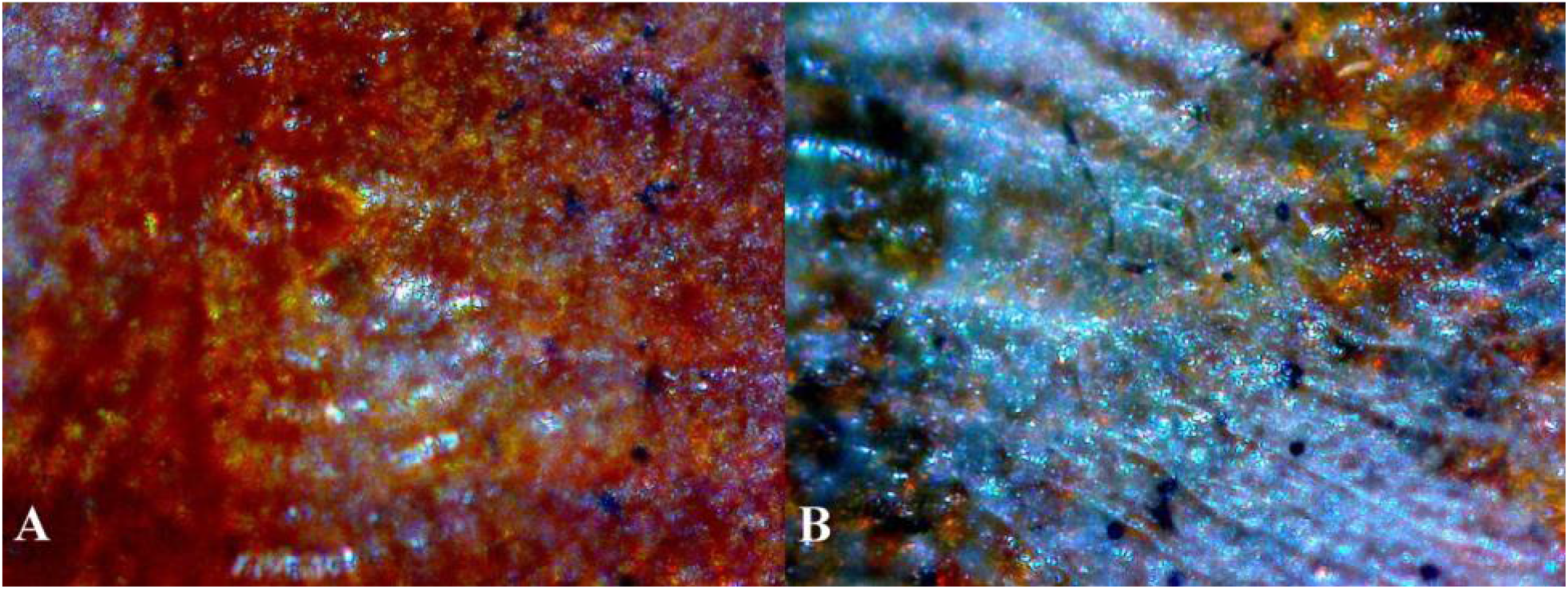
**(A)** 27 Pb 100X 34 *bb, Pb/Pb* (*dissected*) transmitted light revealing increased violet-blue iridophore expression. **(B)** 28 non-Pb 100X 1 *bb, pb/pb* (*dissected*) transmitted light revealing balanced violet-blue iridophores.

#### Rear Peduncle Spot (dissected)

### VII. Cellular Comparison: late coloration in Golden gg, *Pb/pb* vs. Golden non-Pb gg*, pb/pb* male Guppies (*Poecilia reticulata*)

Macroscopically ornamental spot coloration is modified from a highly reflective orange to a “pinkish-purple” in Golden (g, **Fig 30**) (Goodrich 1944). Orange ornament is converted to pinkish-purple in Pb by xanthophore removal, resulting in a smaller area of pigmentation and revealing increased underlying violet-blue structural color. Golden in turn reduces and aggregates melanophores, resulting in even further “constriction” of color pigments as compared to non-Golden.

**Fig 30.**
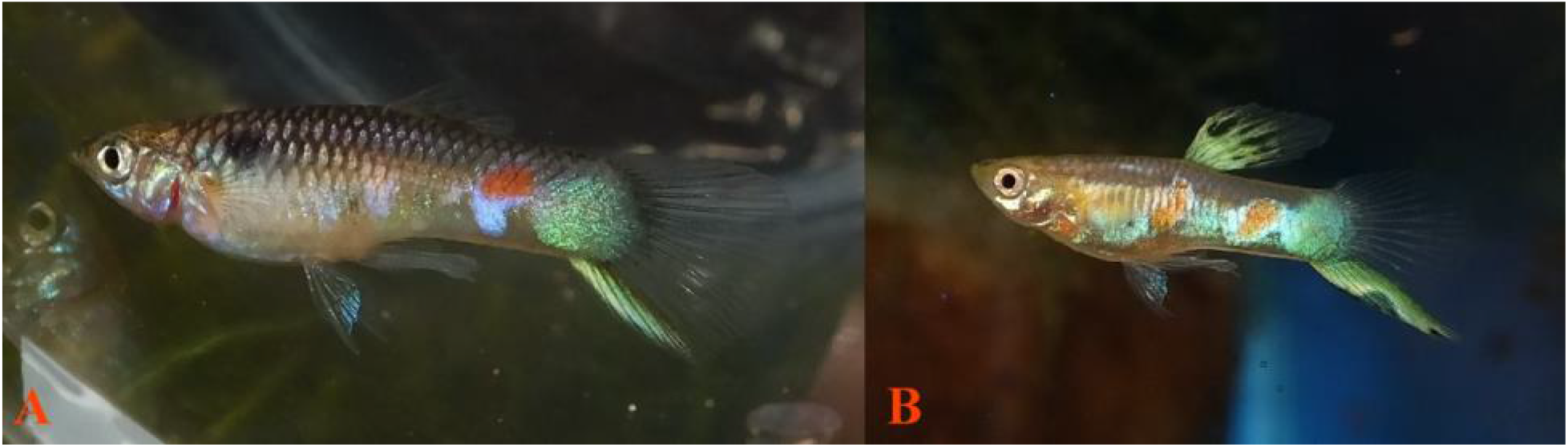
**(A)** Golden Pb *gg, Pb/pb* expressing “constricted” modified pinkish-purple. **(B)** Golden non-Pb gg, *pb/pb* expressing “constricted” orange ornaments.

### VIII. Cellular Comparison: homozygous Pb *Pb/Pb*, heterozygous Pb *Pb/pb* and non-Pb *pb/pb* under reflected, transmitted, reflected and transmitted lighting

Our study revealed that the presence of all major classes of chromatophores (melanophores, xanthophores, erythrophores, violet-blue iridophores) and crystalline platelets were present in Pb and non-Pb condition **(Fig 31-49)**. Collected and clustered xanthophore populations were reduced in heterozygous Pb condition and removed in homozygous Pb condition, as seen macroscopically **(Fig 31, 36, 41** and **44)** and microscopically **(Fig 32-35, 37-40, 42-43** and **45-49)**. The retention of isolated xanthophores, found in all parts of the body and fins in “wild-type”, remained intact in heterozygous and homozygous Pb condition. As previously noted, Pb in itself has little or no effect on erythrophore populations.

**Fig 31.**
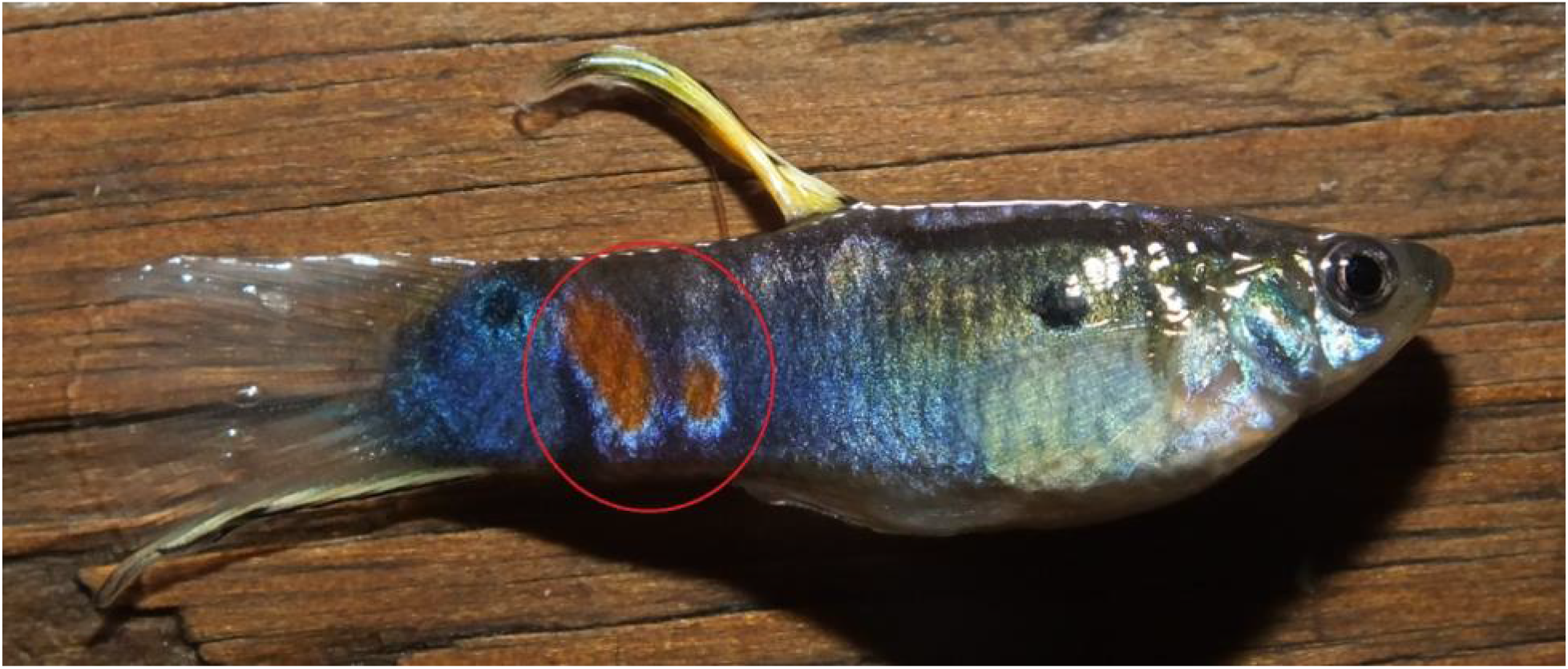
13 Pb male (grey) – Homozygous *Pb/Pb*. All slide images taken from posterior orange spot (red circle).

**Fig 32.**
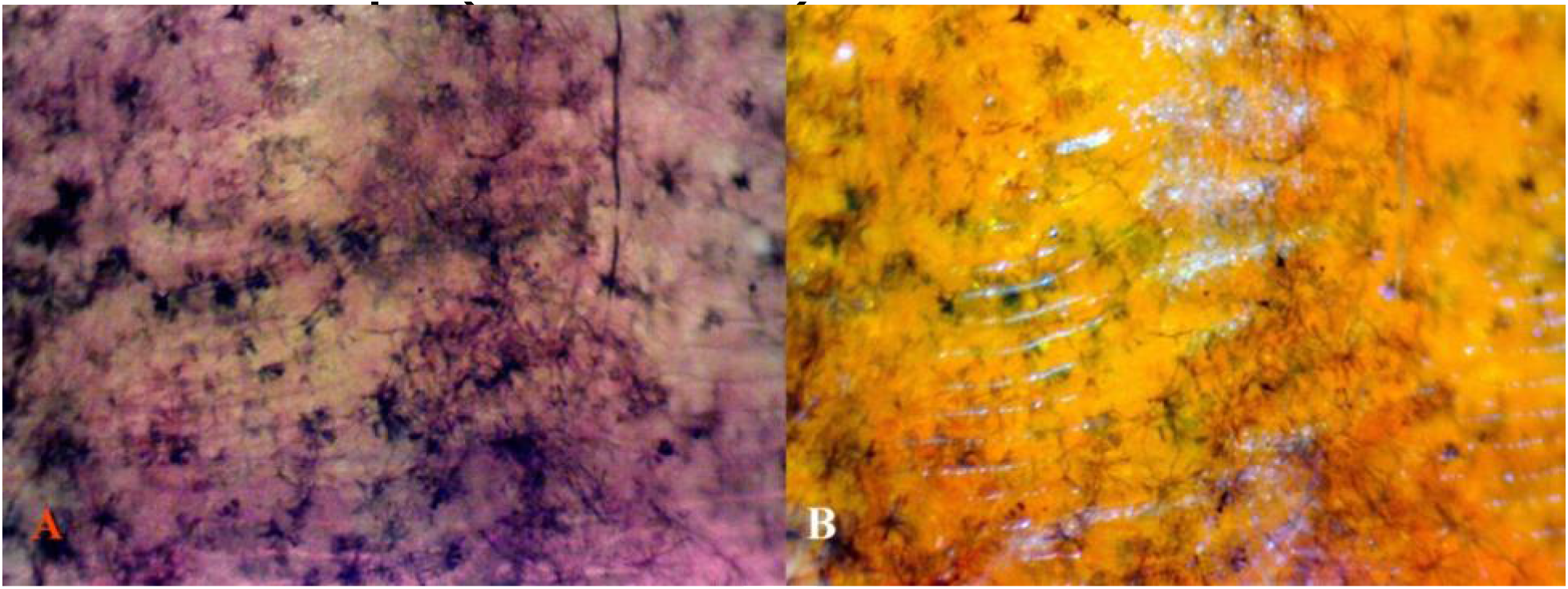
**(A)** 13 Pb 40X 4 *PbPb* transmitted light white balance adjusted. Erythrophore positions are clearly indicated with transmitted light and adjusted white balance. **(B)** The same field, reflected/transmitted light.

**Fig 33.**
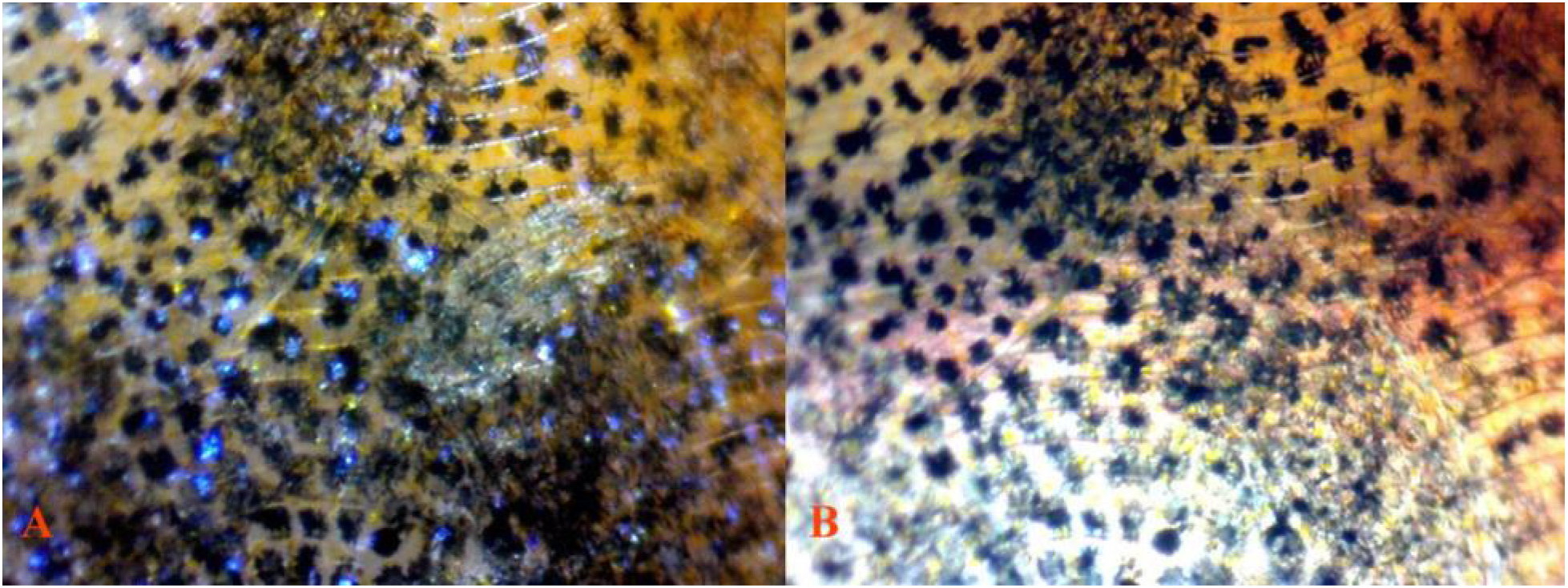
**(A)** 13 Pb 40X 5 *PbP*b reflected light. Reflected light reveals that most of reflective qualities are produced by violet-blue iridophores within scale rings and dendritic melanophore-iridophore chromatophore units. **(B)** The same field, transmitted light white balance adjusted.

**Fig 34.**
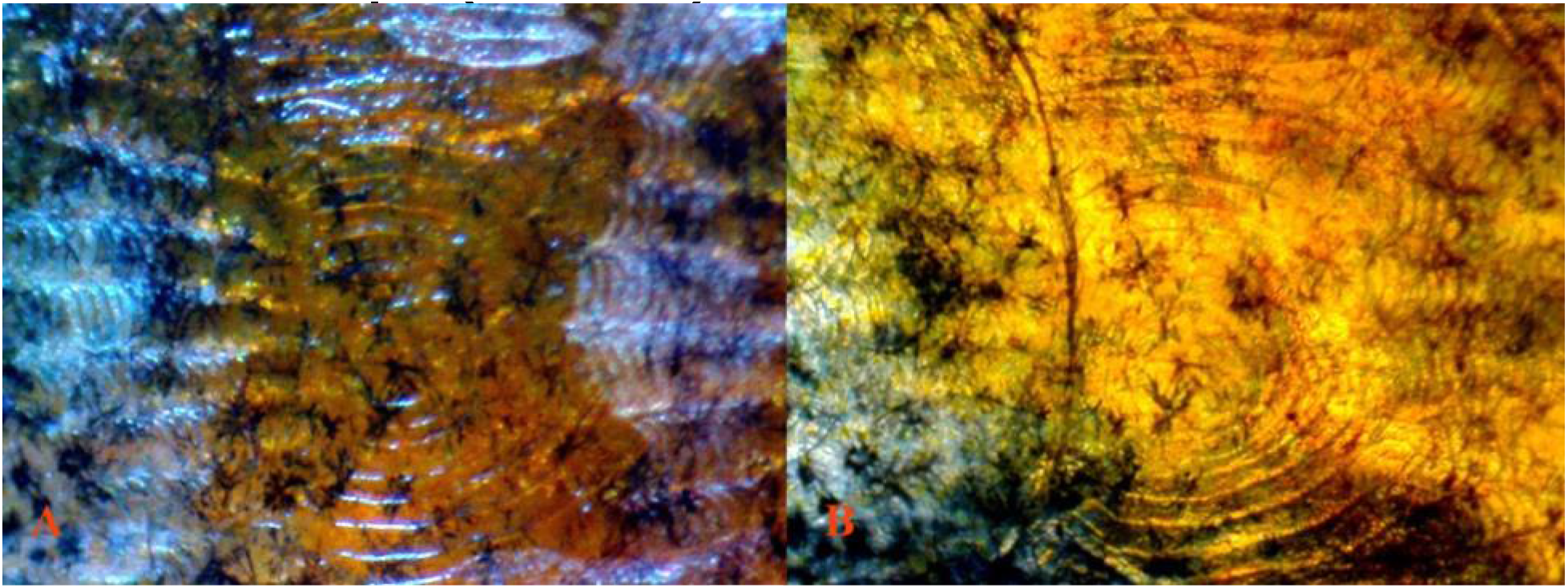
**(A)** 13 Pb 40X 4 *Pb/Pb* reflected light. Violet-blue iridophores shown residing below xantho-erythrophores, and contributing to reflective qualities. The expected presence of “chain-like” dendritic melanophores present along scale edging to form reticulation, dendrites are “extreme” in shape compared to Pb expression. **(B)** The same field, transmitted light.

**Fig 35.**
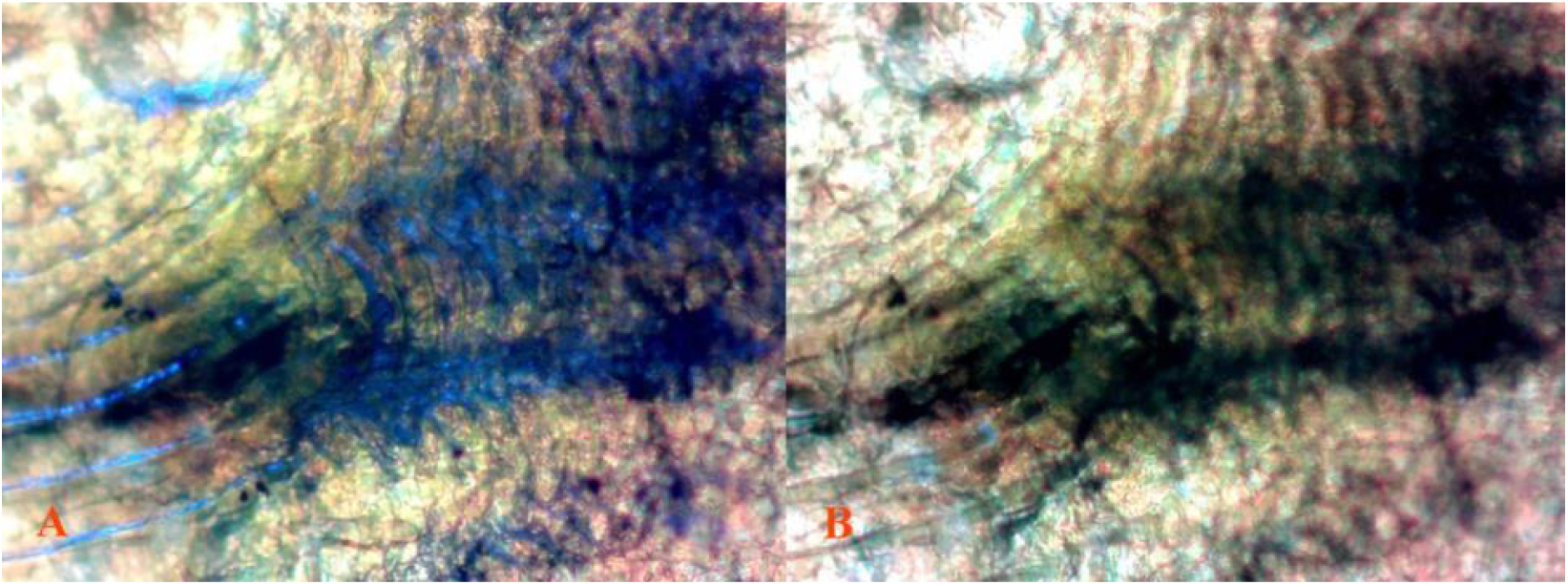
**(A)** 13 Pb 100X 5 *Pb/Pb* reflected/transmitted light white balance adjusted. **(B)** The same field, transmitted light white balance adjusted. In the absence of reflected light iridophore reflective qualities are severely reduced, and as part of dendritic melanophore-iridophore chromatophore units they contribute to dark pattern ornamentation.

**Fig 36.**
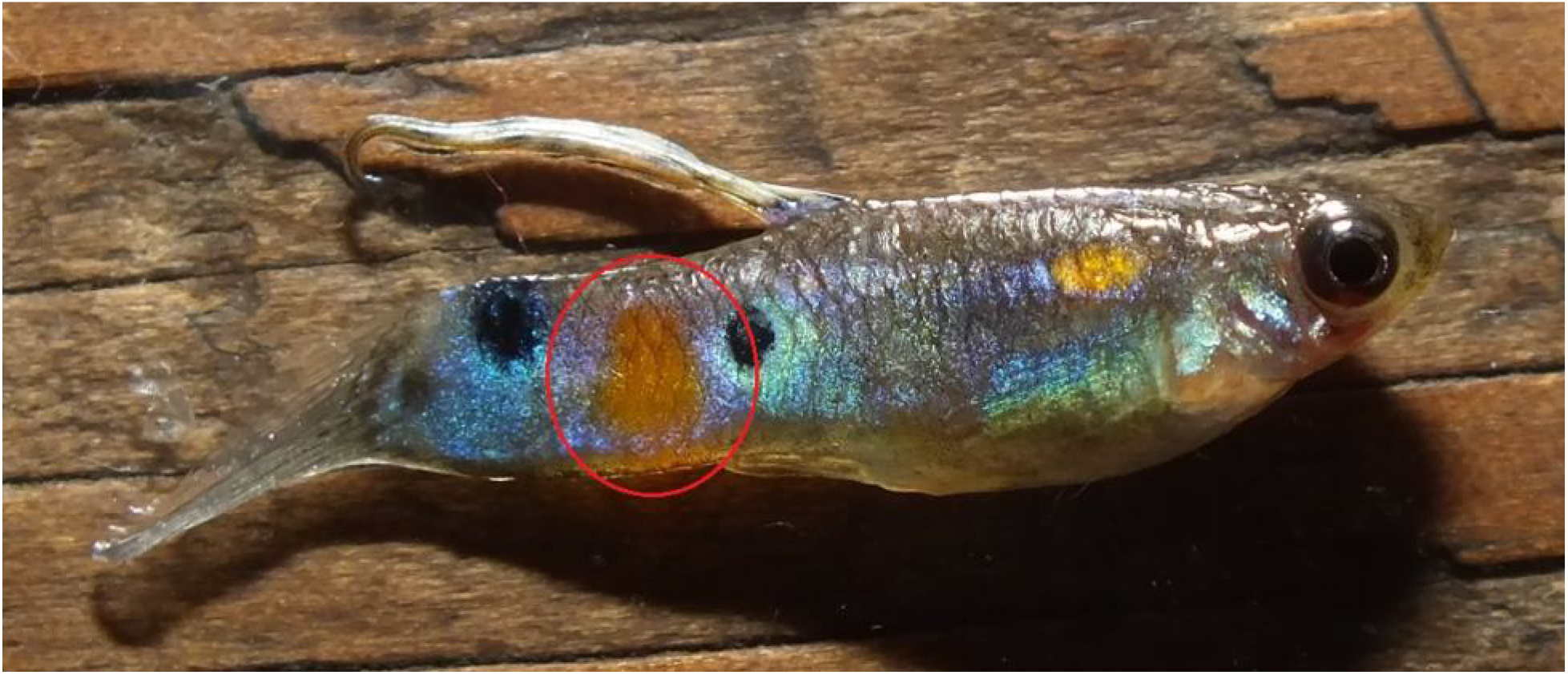
15 Pb male (grey Pingtung feral) – Heterozygous *Pb/pb*. All slide images taken from posterior orange spot (red circle).

**Fig 37.**
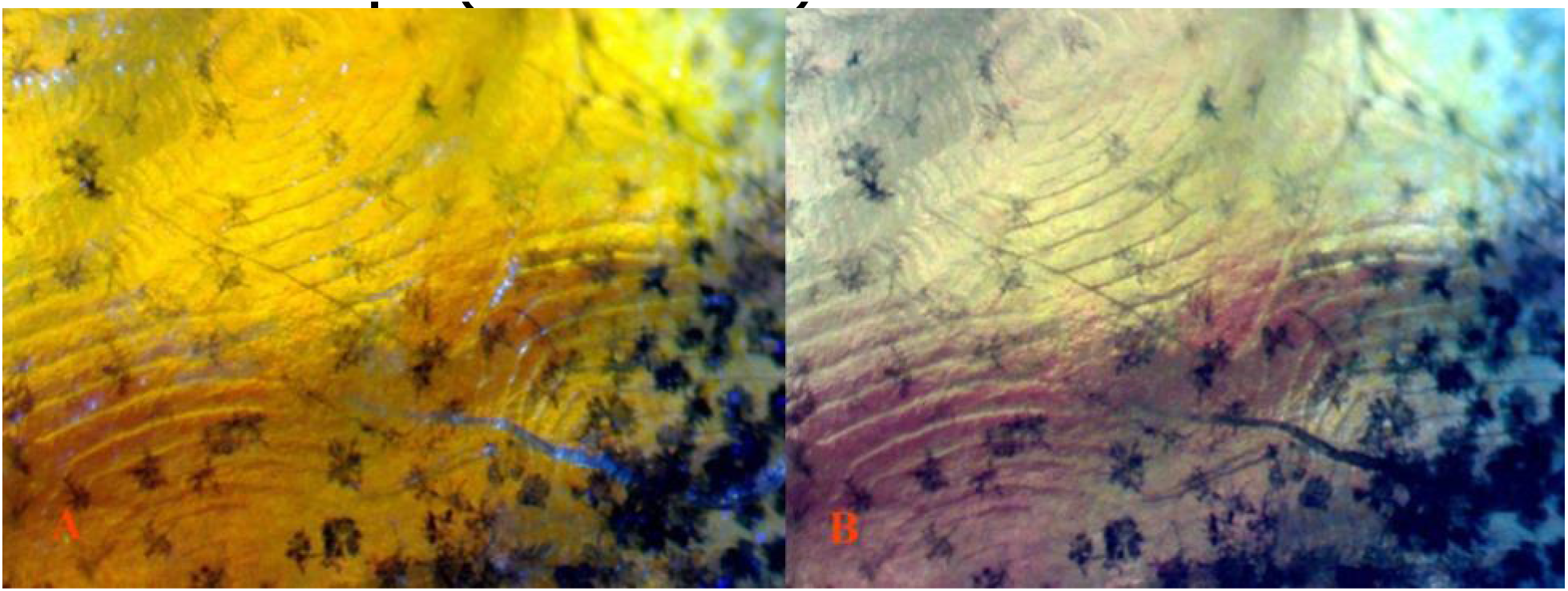
**(A)** 15 Pb 40X 5 *Pb/pb* reflected /transmitted light. The same field, **(B)** transmitted light white balance adjusted. Lacking reflected light xantho-erythrophore expression is reduced, revealing violet-blue iridophore populations near outer edge of spotting ornament.

**Fig 38.**
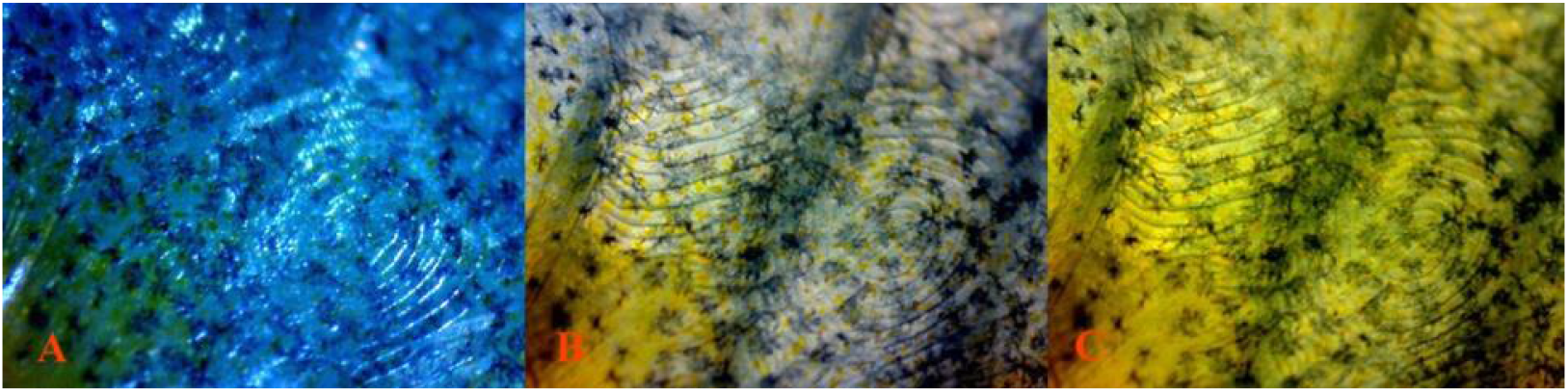
**(A)** 15 Pb 40X 6 *Pb/pb* reflected light. High density lower level and scale ring violet-blue iridophores residing at the edge of spotting ornament. **(B)** The same field, transmitted light white balance adjusted. **(C)** The same field, transmitted. Violet-blue iridophores, within ectopic dendritic melanophore-iridophore chromatophore units, while lacking reflective qualities remain visible under all transmitted light conditions.

**Fig 39.**
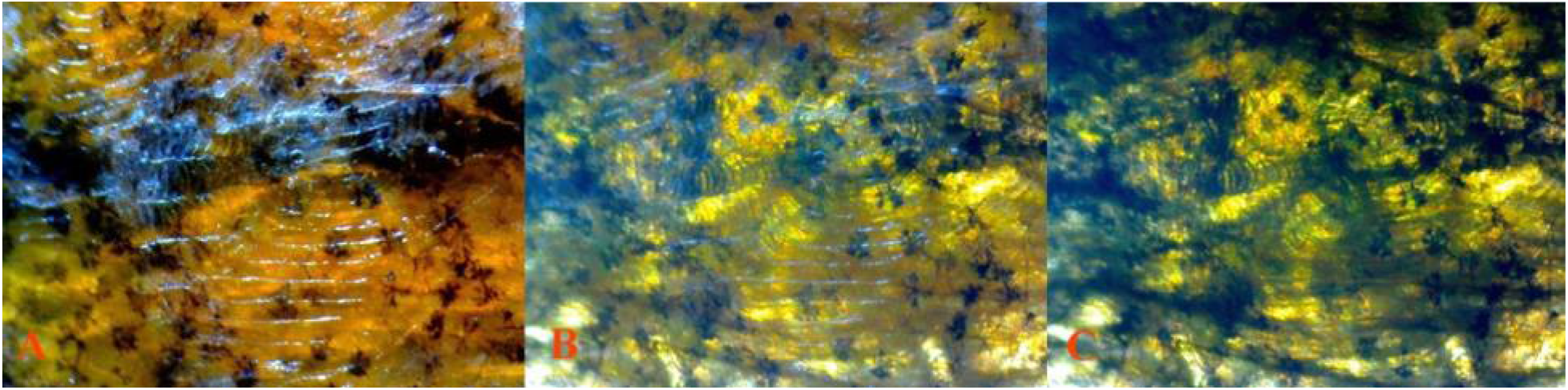
**(A)** 15 Pb 40X 2 *Pb/pb* reflected light. Extreme dendritic melanophores along scale edging. **(B)** The same field, reflected/transmitted light. **(C)** The same field, transmitted. Dendritic melanophore-iridophore chromatophore units are shown to play a major role in dark pattern composition under reduced transmitted light.

**Fig 40.**
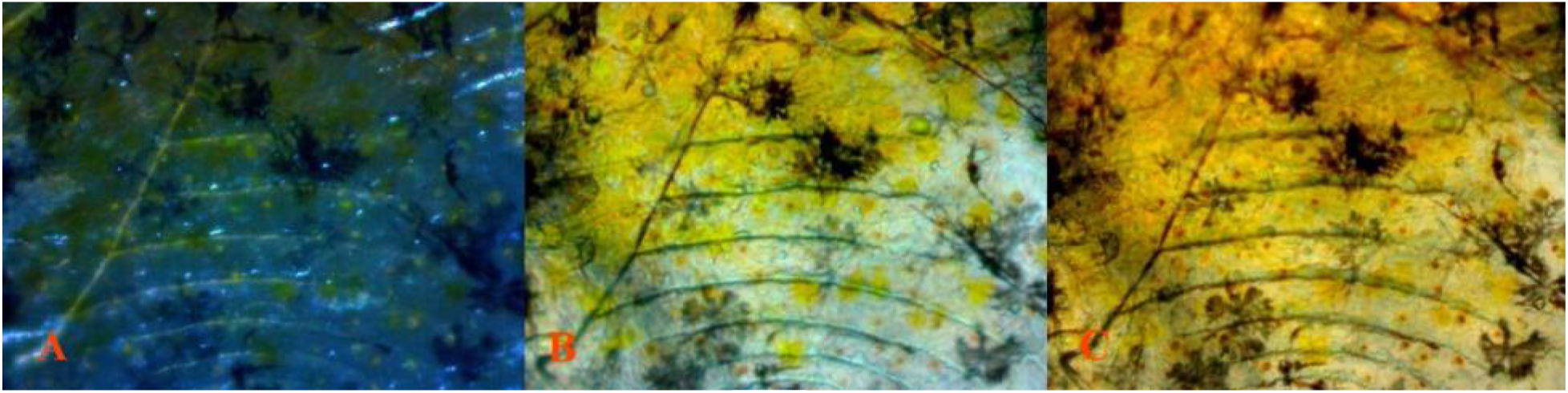
**(A)** 15 Pb 100X 4 *Pb/pb* reflected light. Dendritic xanthophore structures are highlighted under reflected light. **(B)** The same field, reflected/transmitted. Erythrophore expression is minimal under combined reflected/transmitted light near the edge of spotting ornament. **(B)** The same field, transmitted light. Isolated erythrophore expression highlighted under transmitted light.

**Fig 41.**
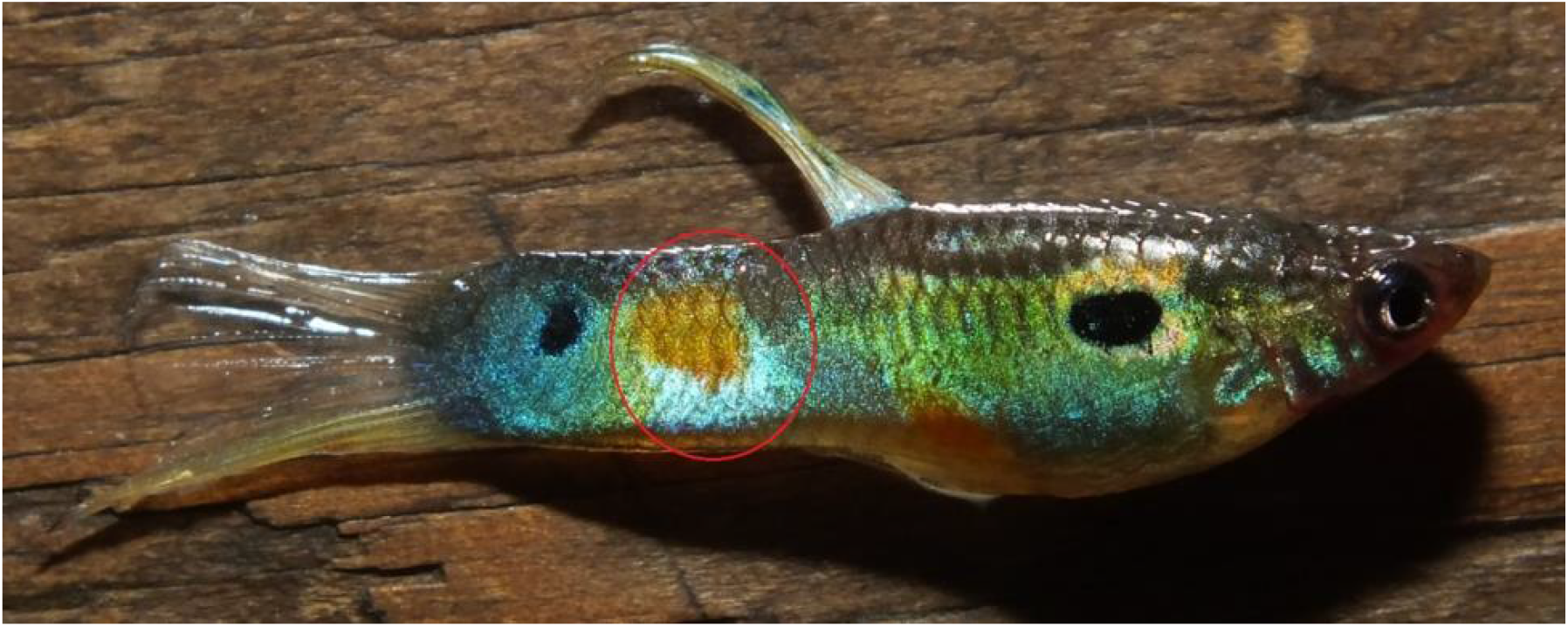
14 non-Pb male (grey) *pb/pb*. All slide images taken from posterior orange spot (red circle).

**Fig 42.**
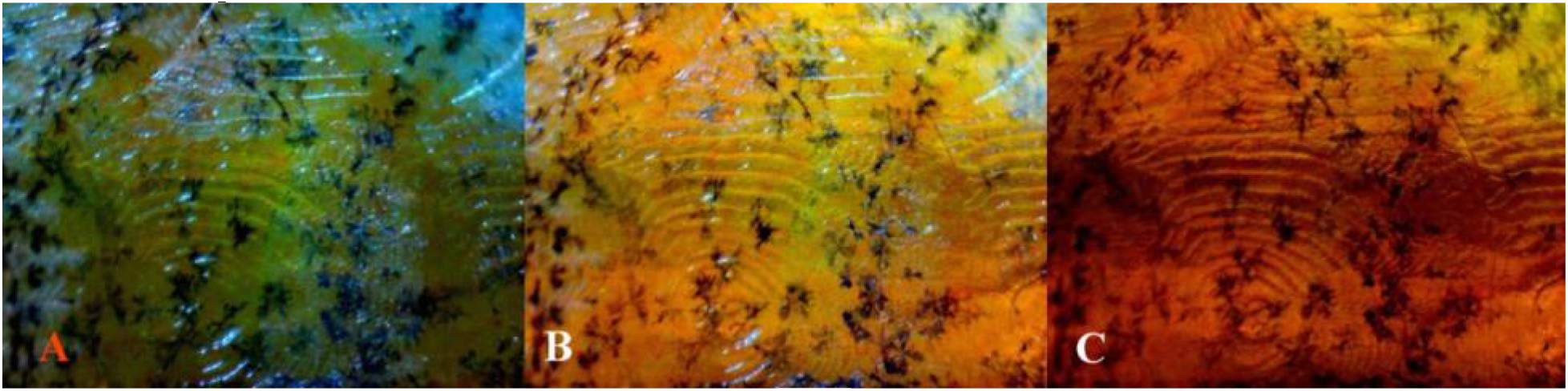
**(A)** 14 non-Pb 40X 1 *pb/pb* reflected light. Location of violet-blue iridophores is shown to be not limited to non-color pigmented areas. Indicating a solid structural color “layer” regardless of presence or absence of color pigments. **(B)** The same field, reflected/transmitted light. Combined reflected/transmitted light suggestive of positioning of carotenoid orange pigment to be slightly above that of xanthophores. **(C)** The same field, transmitted light. Transmitted light again allows erythrophore expression to “overpwer” that of xanthophores.

**Fig 43.**
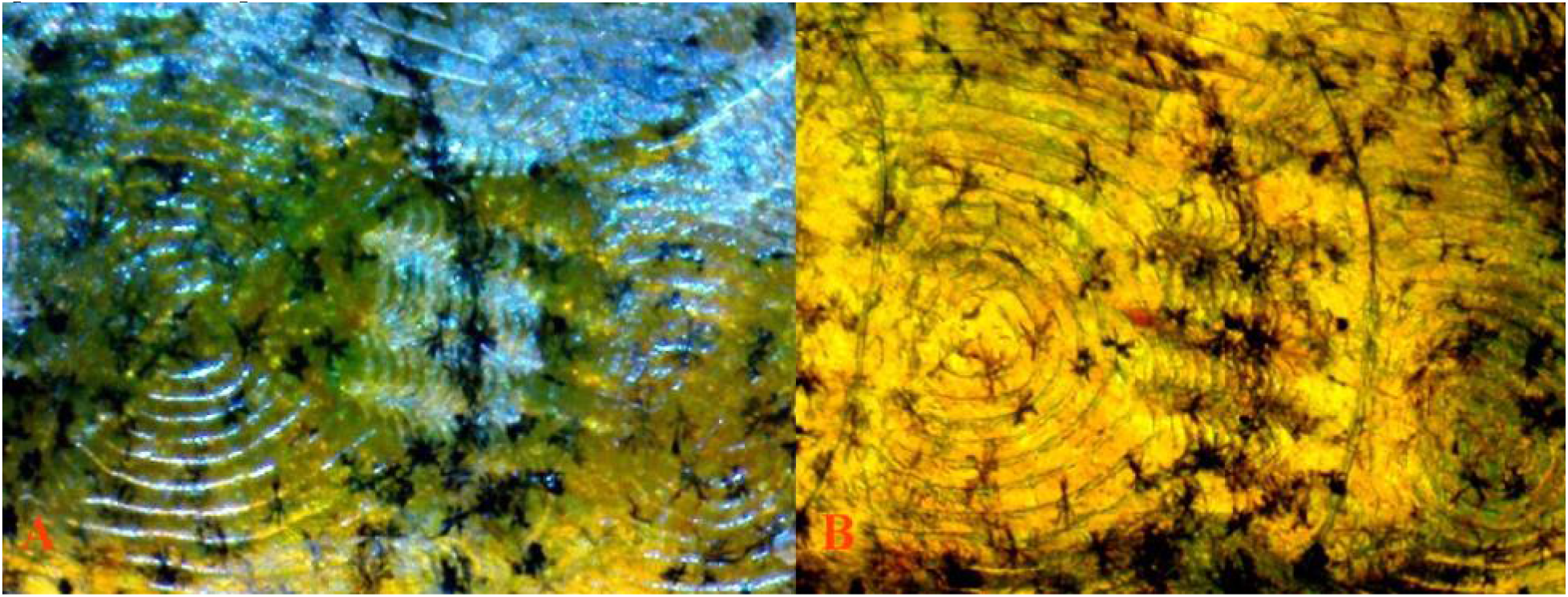
**(A)** 14 non-Pb 40X 3 *pb/pb* reflected. Expected presence of “chain-like” dendritic melanophores present along scale edging to form reticulation, though dendrites are not as “extreme” in shape compared to Pb expression. **(B)** The same field, transmitted light. Higher levels of xanthophore expressed in non-Pb

**Fig 44.**
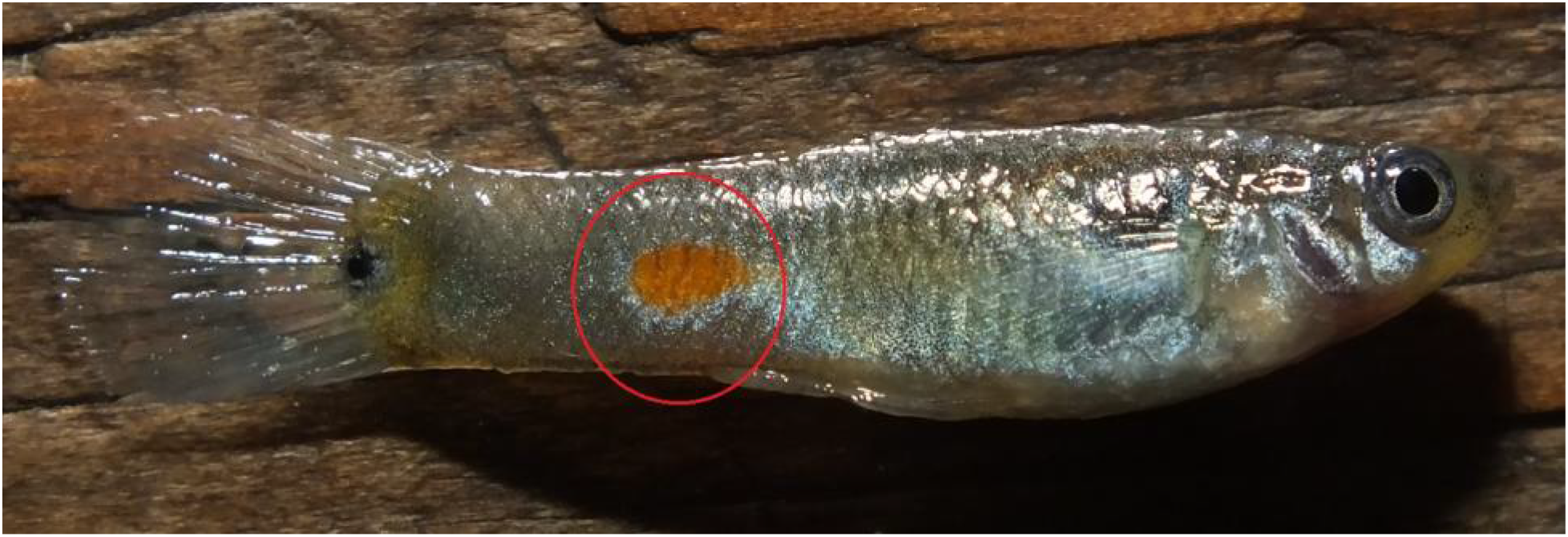
16 non-Pb male (grey Kelly) *pb/pb*. All slide images taken from posterior orange spot (red circle). Reticulation expression is reduced in absence of multiple color ornaments.

**Fig 45.**
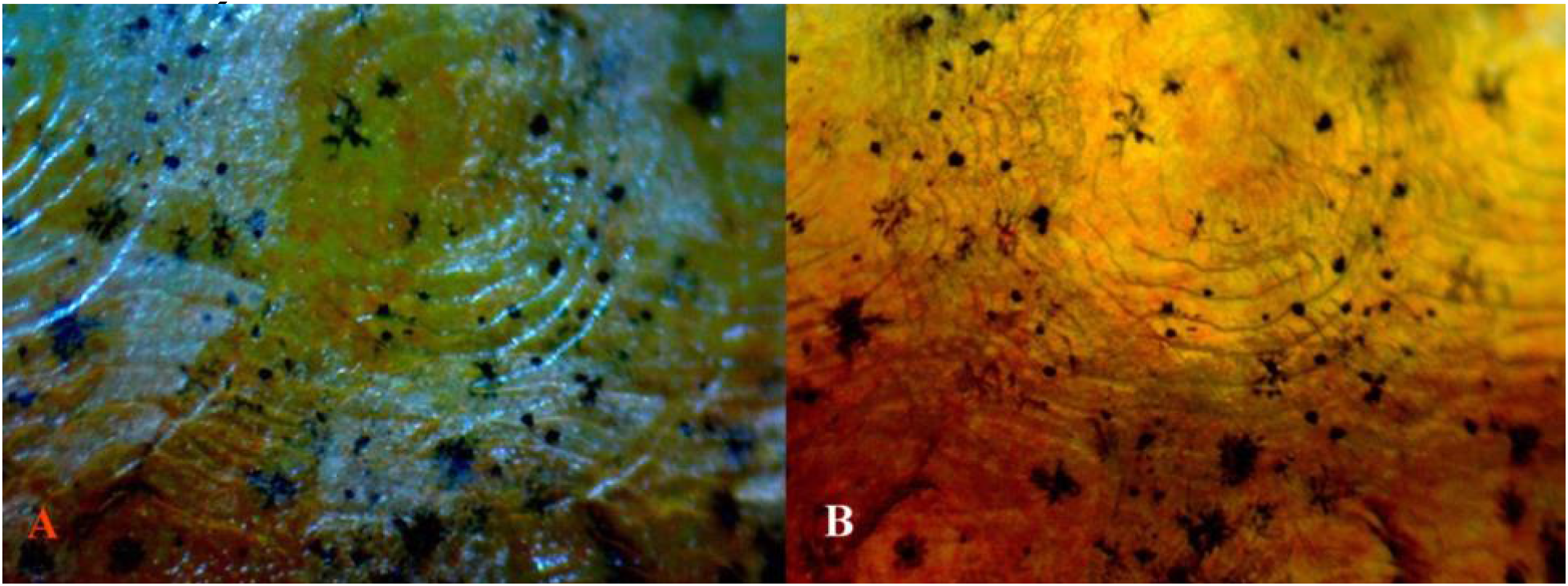
**(A)** 16 non-Pb 40X 2 *pb/pb* reflected light. High concentration of violet-blue iridophores underlying color pigments, in absence of color pigments and within scale rings. **(B)** The same field, transmitted light. Melanophores comprised of punctate, corolla and dendritic cells.

**Fig 46.**
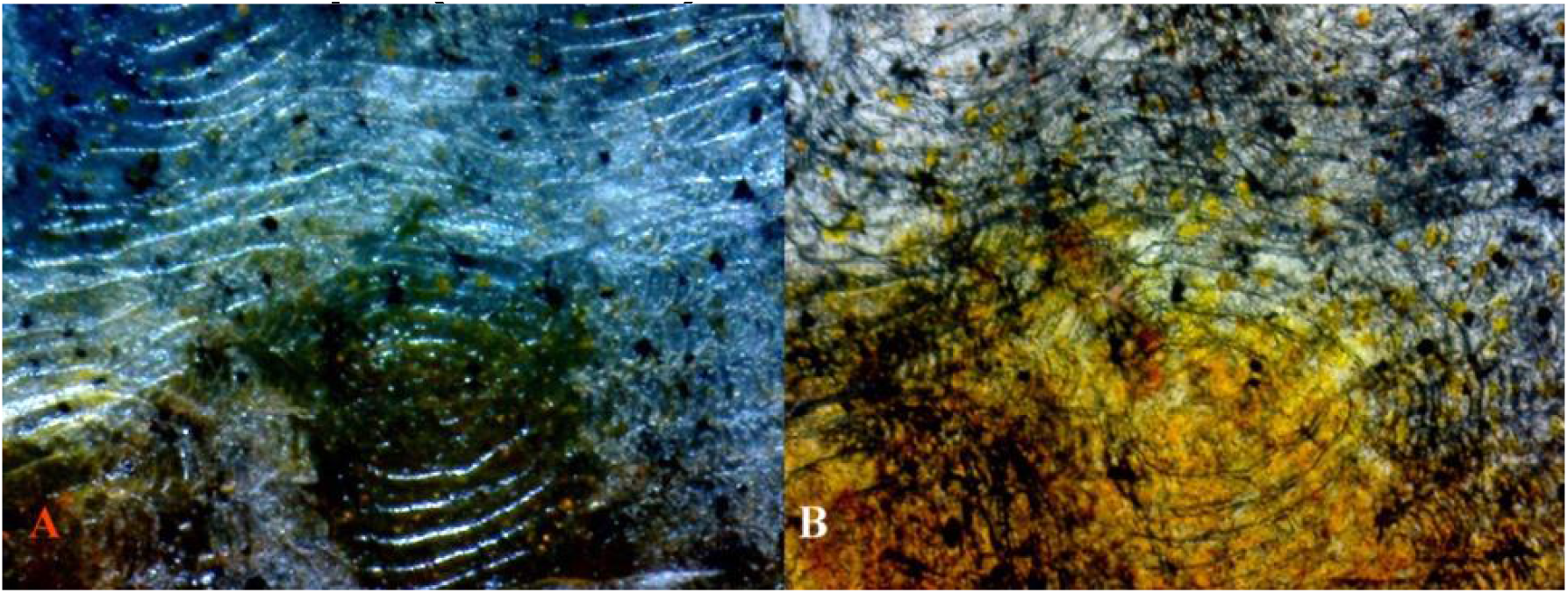
**(A)** 16 non-Pb 40X 2 *pb/pb* reflected light. **(B)** The same field, reflected white balance adjusted. Dendritic melanophore-iridophore chromatophore units comprise much of dark pattern. Erythrophores are visible under transmitted light.

**Fig 47.**
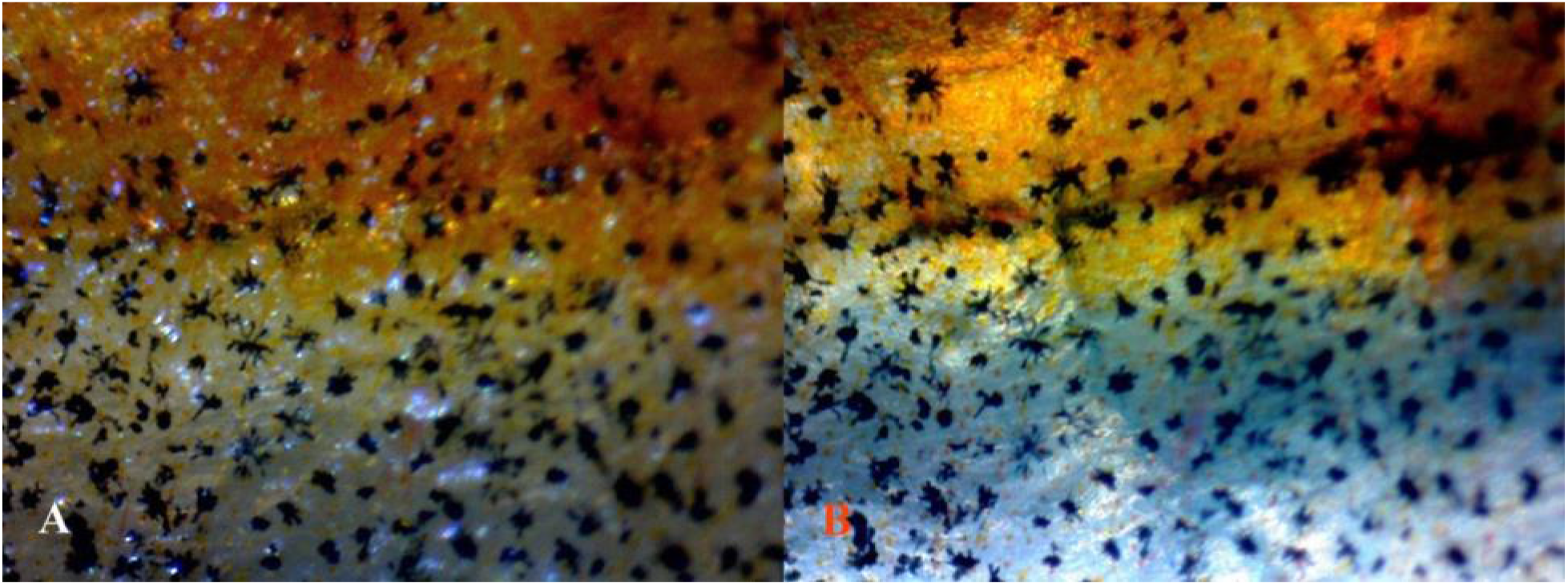
**(A)** 16 non-Pb 40X 6 *pb/pb* reflected light. **(B)** The same field, reflected/transmitted light. This mature male lacks additional spotting ornaments. As a result melanophores appear “less organized” in comparison to heavily ornamented individuals.

**Fig 48.**
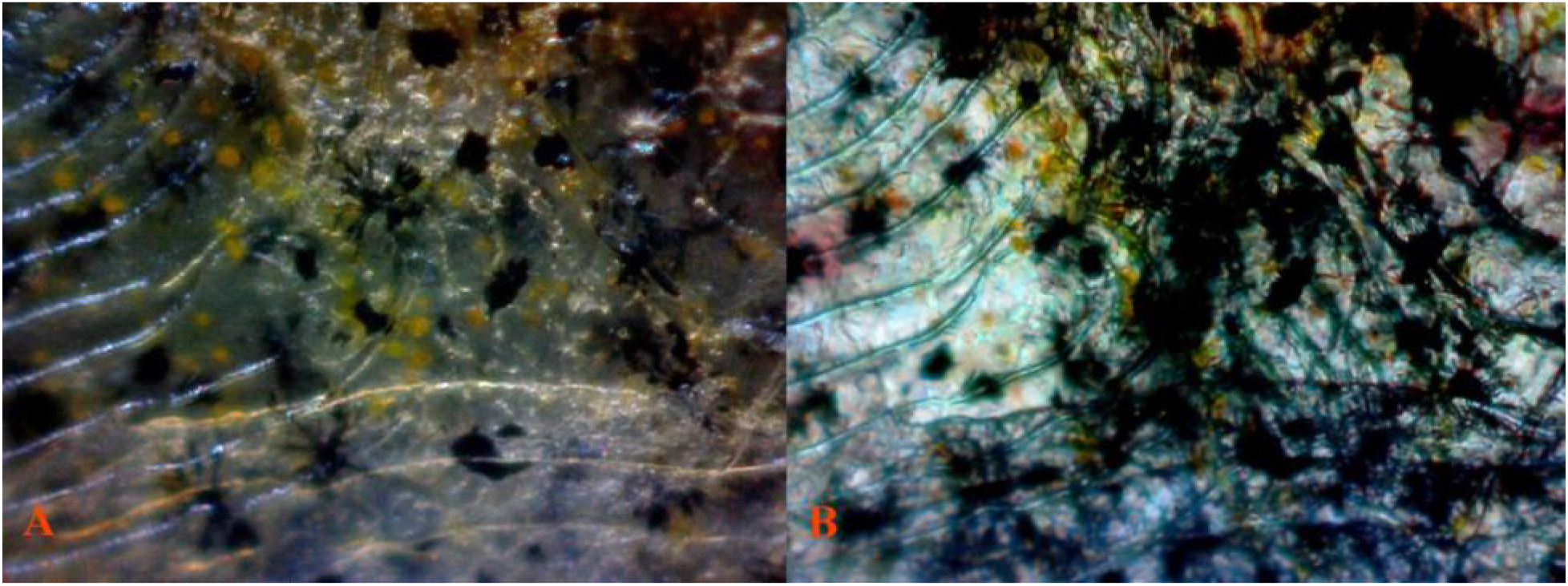
**(A)** 16 non-Pb 100X 2 *pb/pb* reflected light. **(B)** The same field, reflected/transmitted light white balance adjusted. Violet-blue iridophores visible in scale rings under reflected light contributing to reflective qualities. Their presence was also revealed in previous scale dissections. Dendrites under reflected/transmitted light appear much “denser” from interactions with violet-blue iridophores. Variation in dendrite shape at a single focal length indicates an ectopic orientation at distinct layers.

**Fig 49.**
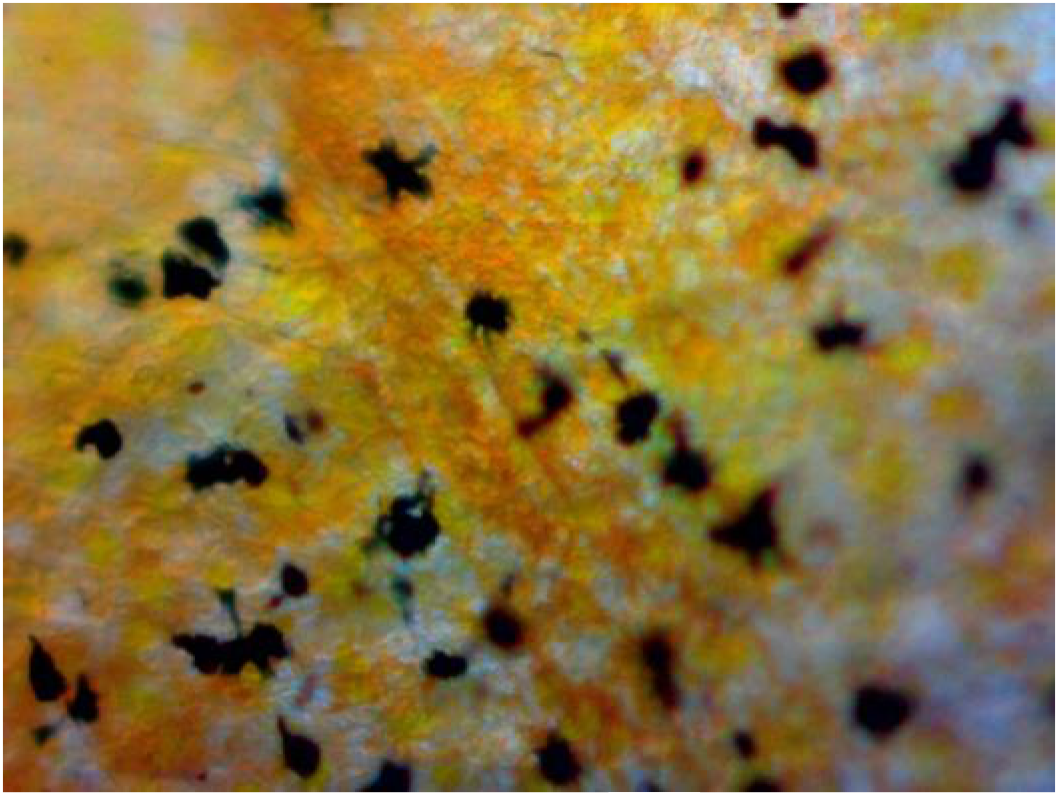
**(A)** 16 non-Pb 100X 9 *pb/pb* transmitted light. Xantho-erythrophores appear “balanced” in expression lacking Pb modification. No extreme modification of dendrites is visible. Well defined layers between structural color and color pigment. Even with minimal dendritic extension, some collection of violet-blue iridophores into ectopic dendritic melanophore-iridophore chromatophore units is still visible.

The presence of underlying iridophores, lacking a xanthophore (yellow) overlay in Pb, with retained erythrophore (orange) overly produces a distinct “pinkish-purple” coloration in heterozygous and homozygous Pb condition, which is visible both macroscopically and microscopically. The alteration of orange spotting is complete in homozygous Pb, and limited to specific regions in heterozygous Pb (*See:* Bias and Squire 2017a).

Homozygous Pb specimens **(Fig 31-35)** exhibited an “overall” higher incidence of violet iridophores, a proliferation of dendritic melanophores and a dense layer of violet-blue iridophores, as compared to heterozygous Pb **(Fig 36-40)** and non-Pb **(Fig 41-49)**. A “darker” appearance in Pb vs. non-Pb appears to largely be the result of modification of existing melanophore structures (corolla and punctate) into extended dendrites. An increase in melanophore population levels is not evidenced.

Dendritic melanophores in heterozygous and homozygous Pb condition, for mature individuals, reveal that dendrites are extremely extended and finer in appearance. Dendrites are linked together in “chain-like” strings intermingled with violet-blue iridophores in chromatophore units forming the reticulated pattern and in isolated units. This chain-like expression is minimally present in non-Pb **(Fig 42-43, 45-49)**, further present in heterozygous Pb **(Fig 37-40)**, and amplified in homozygous Pb **(Fig 32-35)**. Within the rear peduncle orange spot and surrounding edges a noticeable absence of corolla and punctate melanophores is often evident in Pb. This absence was reduced in other regions of the body or was specific to individuals.

#### Rear Peduncle Spot (non-dissected)

#### Rear Peduncle Spot (dissected)

#### Rear Peduncle Spot (non-dissected)

#### Rear Peduncle Spot (dissected)

#### Rear Peduncle Spot (non-dissected)

#### Rear Peduncle Spot (dissected)

#### Rear Peduncle Spot (non-dissected)

#### Rear Peduncle Spot (dissected)

### IX. Subcutaneous and Spinal Chromatophores

Macroscopic qualities of non-Pb study male **(Fig 50)**. Melanophore, xanthophore and violet-blue iridophore were found adhering to the spinal column and ribs **(Fig 51A-B**). No evidence of subcutaneous erythrophores was detected below the dermis, though suspected in micro-dissected the xanthophore cluster **(Fig 52)**; all “red coloration” identified as blood cells. This raises an issue in regard to the synthesized pteridine and dietary carotenoid pigment resource allocation in non-visible locations (Goodwin 1984; Grether 1999], 2001). Subcutaneous melanophore, xanthophore violet-blue iridophore, leucophore and crystalline platelets **(Fig 53-54)** were found resident in low numbers in dissected deep tissue. Their presence at these locations is well below the dermis (Bagnara 1968).

**Fig 50.**
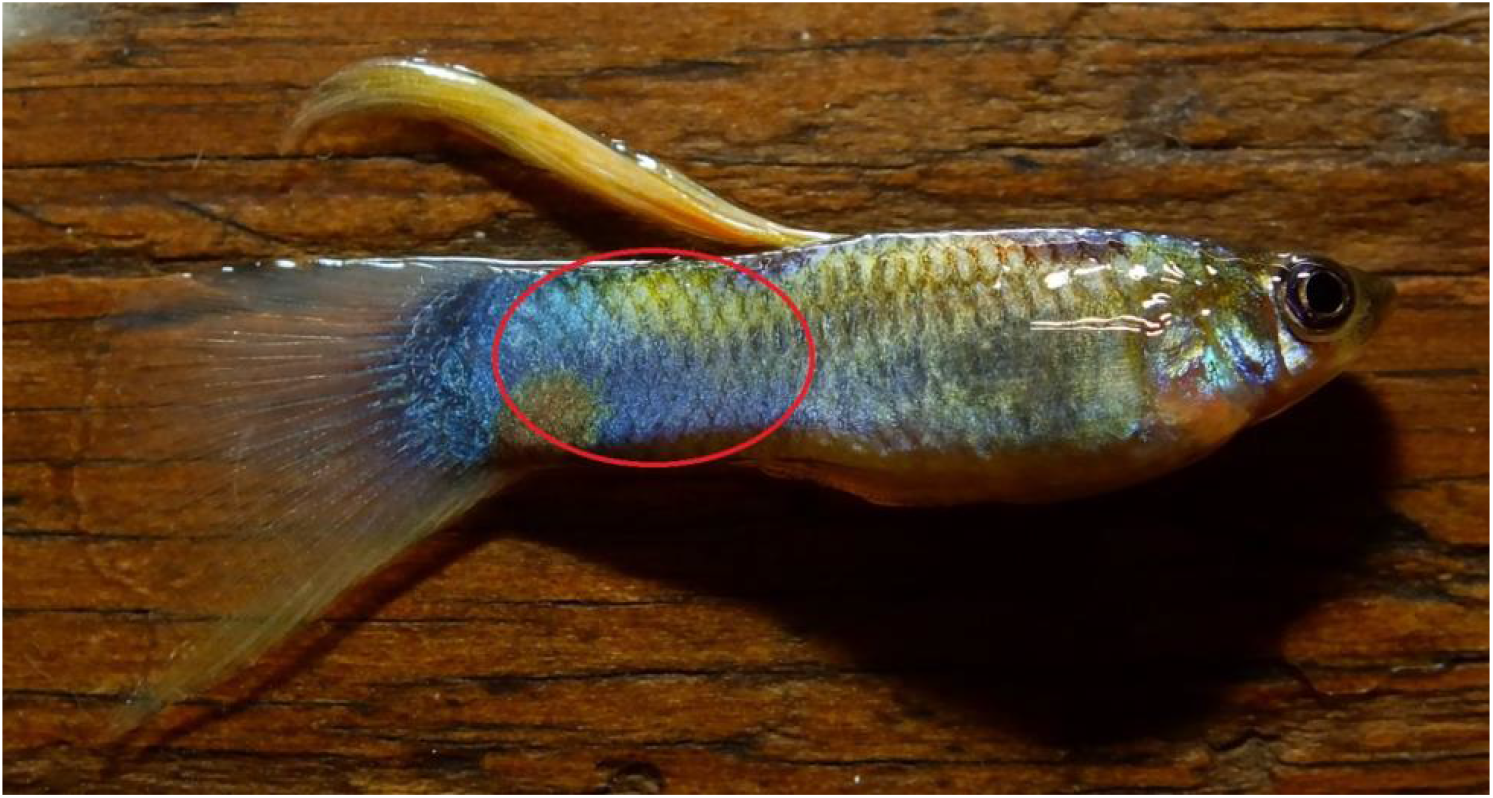
25 non-Pb (McWhite) *pb/pb*. All slide images taken from posterior peduncle (red circle).

**Fig 51.**
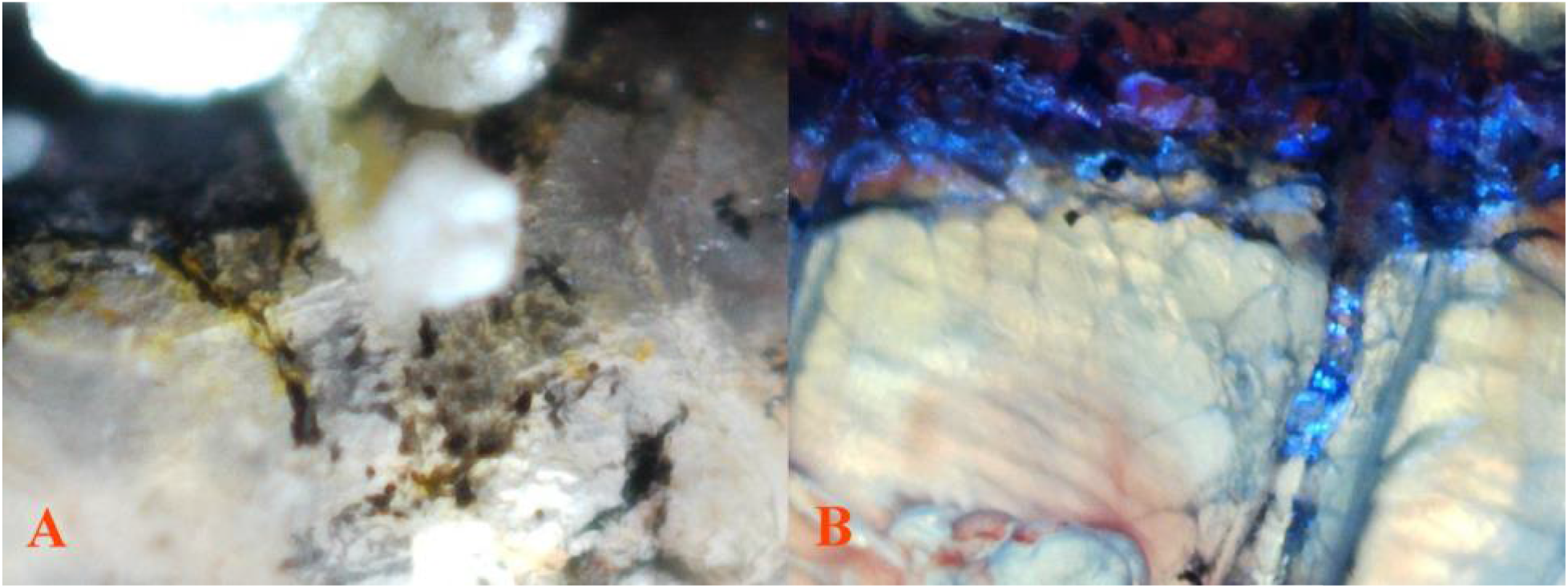
**(A)** 25 non-Pb 100X 9 (*micro-dissected spinal column*) reflected light. Visible dendritic melanophores and xanthophores residing on the freshly extracted and cleaned spinal column, rinsed with saline solution. **(B)** The same fish (*partially dissected spinal column*) reflected/transmitted light. Visible violet-blue iridophores are residing just above dendritic melanophores and xanthophores on the freshly dissected and cleaned spinal column, rinsed with saline solution.

**Fig 52.**
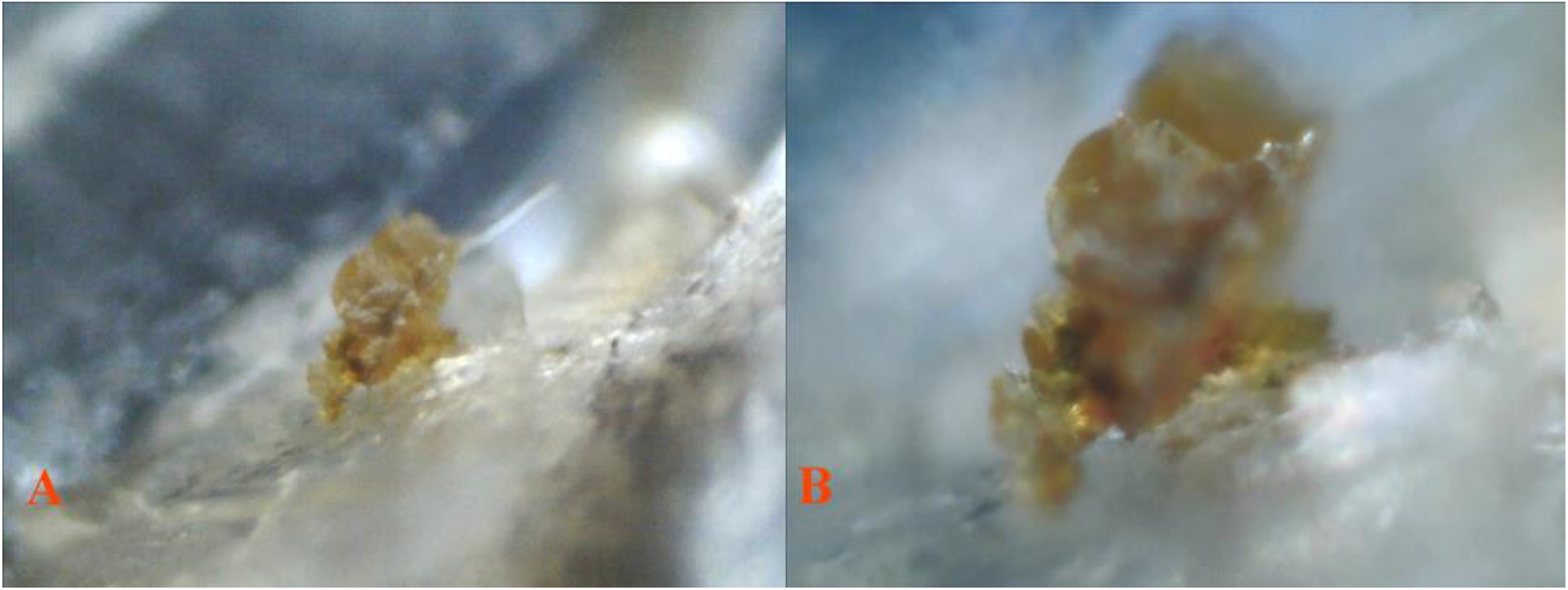
**(A)** 25 non-Pb 40X 1 (*micro-dissected tissue*) reflected light. **(B)** The same field 100X 1 (*micro-dissected tissue*) reflected light. Isolated yellow xanthophore cluster and reflective crystalline platelets from subcutaneous deep tissue sample.

**Fig 53.**
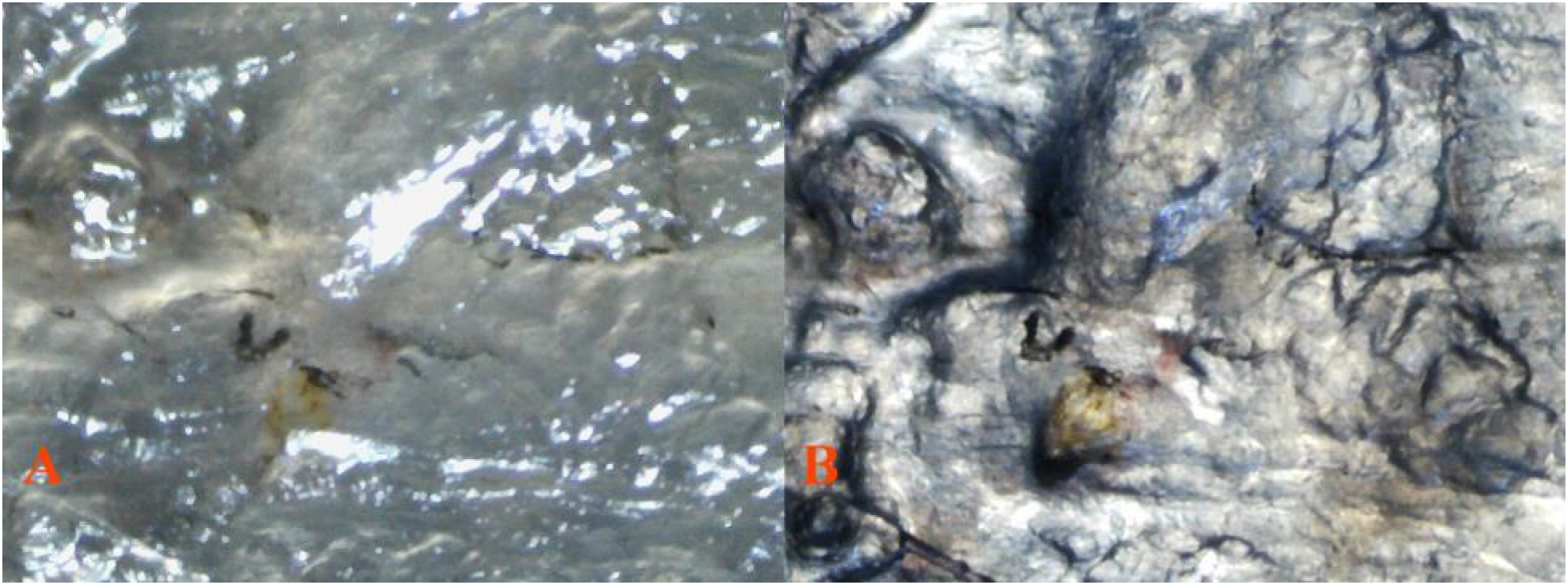
Proximal spine side up (*inverted*) views of same location. **(A)** 25 non-Pb 40X 1 (*dissected tissue*) reflected light. Visibly radiating dendritic melanophores and xanthophores are present. Bright white areas are now inverted crystalline platelets above scattered violet-blue iridophores. This shows that crystals are reflective both distally and proximally under any available reflected light. White leucophores are “grey” in appearance. **(B)** The same field 40X 1 (*dissected tissue*) reflected/transmitted light. Aggregated dendritic melanophores and violet-blue iridophores forming ectopic melanophore-iridophore chromatophore units are visible with isolated yellow xanthophore cluster. Reflective crystalline platelets remain white. White leucophores are “grey” in appearance.

**Fig 54.**
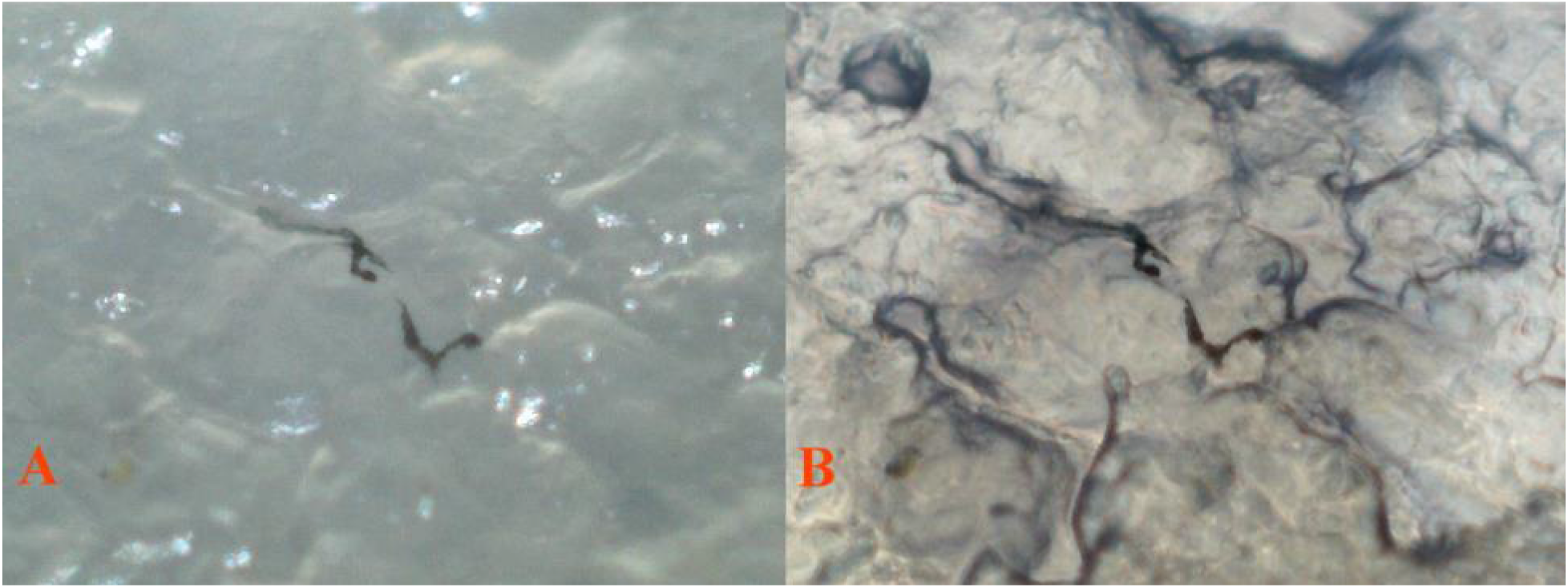
Proximal spine side up (*inverted*) views of same location. **(A)** 25 non-Pb 100X 1 (*dissected tissue*) reflected light. Visibly radiating dendritic melanophores are present. Bright white areas are now inverted crystalline platelets above scattered violet-blue iridophores. Showing that crystals are reflective both distal and proximal under any available reflected light. White leucophores are “grey” in appearance. **(B)** The same field, (*dissected tissue*) reflected/transmitted light. Aggregated dendritic melanophores and violet-blue iridophores forming ectopic melanophore-iridophore chromatophore units are visible. Reflective crystalline platelets remain white. White leucophores are “grey” in appearance.

## Discussion and Conclusions

Our study revealed all major classes of chromatophores (melanophores, xanthophores, erythrophores, and violet-blue iridophores) and also crystalline platelets were present in both Pb and non-Pb dermal layers. All were found present in and around the eye structure. Scales contained their own distinct populations, and all but erythrophores were found. In dermal layers, the dendrites of ectopic dendritic melanophore structures appeared to be tilted when melanosomes are fully dispersed. Collected and clustered xanthophore populations were reduced in heterozygous Pb condition and nearly removed in homozygous condition. Isolated clustered xanthophore found over all parts of the body and fins in “wild-type”, remained intact in both heterozygous and homozygous Pb condition.

Melanophores, xanthophores, violet-blue iridophores, leucophores and crystalline platelets were found resident in subcutaneous deep tissue samples well below the dermis. Deep tissue crystalline platelet orientation appears to be tilted, as they are reflective both distally and proximally under reflected light. Dendritic melanophores radiate in deep tissue indicating 3-dimensional (3-D) orientation within the subcutaneous tissue. Melanophores, xanthophores, violet-blue iridophores, and also crystalline platelets were found adhering to the spinal column.

An aggregation of dendritic melanophores and violet-blue iridophores forming ectopic melanophore-iridophore chromatophore units (*Fig. 1 and Fig. 6c*, Kottler 2014; Fig 17A-B, Bias and Squire 2017a), in cutaneous and subcutaneous levels is supportive of structural color cells residing at multiple layers, and being above and below the distinct iridophore layer as described in the Bagnara Dermal Chromatophore Unit (Bagnara 1968).

The presence of underlying dermal level violet-blue iridophores and crystalline platelets, lacking a xanthophore (yellow) overlay in Pb, with a retained erythrophore (orange) layer produces a distinct “pinkish-purple” coloration in heterozygous and homozygous Pb condition, which is visible both macroscopically and microscopically. An increase in reflective qualities of violet-blue structural pigment is evidenced in Pb. Based on our study of “specific phenotypes”, Pb expresses a higher amount of violet iridophores or more balanced ratio of violet-blue iridophores. Violet-blue iridophores are shown, in both Pb and non-Pb, to form evenly distributed layers in the presence or absence of color pigments. There is evidence that xanthophores and erythrophores may reside at slightly diverse layers or angles within colored ornaments.

The violet-blue iridophore chromatophore unit (Fig 2A-B, Bias and Squire 2017a) and the removal of xanthophores by Pb modification is required to produce an all-purple phenotype. The Purple gene has the ability to modify existing genome-wide chromatophore populations in heterozygous and homozygous condition, with increased visibility in the UV and/or near-UV spectrum. As a result, this demonstrates selection favoring “private” short wavelength signaling (Endler 1991; Millar 2012).

Visually, ornamental spot coloration is modified from a highly reflective orange to a “pinkish-purple” in Grey (g), Blond (b) and Golden (g) (Goodrich 1944). By further removal of erythrophores in European Blau (r) (Dzwillo 1959) and Asian Blau (Ab) (*Undescribed* - see Bias 2015) variants, ornaments are modified to a “violet-blue” revealing the remaining structural color. Pb modification is often most vividly noticeable in grey, blond and albino (Bias and Squire 2017d, *forthcoming*, PB Expression in Domestic Phenotypes).

Microscopically, our results often show minimal structural differentiation between xantho-erythrophores, with differences in population levels, yellow-orange coloration, and collection or clustering of xanthophores. Yellow color cell populations consisting of isolated “wild-type” single cell xanthophores remain intact in conjunction with Pb modification. The same distinctions between heterozygous and homozygous Pb are generally confirmed in macroscopic observations.

The population of melanophores does not appear to drastically increase with Pb modification, only the size and shape of the melanophores themselves are altered. When comparing the macroscopic and microscopic results, between documented autosomal genes modifying melanophores, it becomes apparent that while melanophores are modified by Pb, their modification in size or shape by Blond or Golden is not required for “Purple / Violet” or “pinkish-purple” expression by Pb. Nor is it prevented by the lack of melanophores in Albino (a) (Haskins & Haskins 1948), see examples in: (Bias and Squire 2017d, *forthcoming*, PB Expression in Domestic Phenotypes). Rather, melanophore presence, modification or absence only determines the actual “shade” of pinkish-purple modification in ornamental spots and overall body color when color pigment is present.

Motility of melanophore structures was not addressed in this study, though constriction of melanophores is known to occur (Nayudu 1979). Melanophore constriction was noticed during ocular study as the length of time that each slide sample was observed progressed. Frequent evidence of dendritic and/or motile yellow color pigment (xanthophore) structures was detected in this study. Both when length of time increased between preparations of euthanized specimens, and as the length of time each slide sample was observed progressed. For these reasons preparation was done immediately after euthanasia and observations kept to under an hour. No constriction was found for dendritic and/or motile iridophores, outside of violet-blue iridophore clustering associated with ectopic dendritic melanophores. It was noted on multiple occasions after extended periods of observation that what appeared to iridophores would randomly “fire” during cell death. Yet, after drying and rehydration reflective qualities of iridophores persevere.

Zygosity and genotype specific, the ratio of violet to blue iridophores appears higher in Pb vs. non-Pb. Whether there is an actual increase in iridophore population numbers or simply an increase in visibility, due to reductions or removal of xanthophores and/or altered melanophores was not addressed in this study. Nor was the issue of increased reflectivity in Pb through visibility and possible modification to angles at which crystalline platelets reside beneath iridophore layers and basal level melanophores. Clearly much room exists for further and more complex cellular level research involving Purple Body modification.

Taken as a whole, macroscopic and microscopic results reveal a complex interaction between all major chromatophores types and crystalline platelets is required to produce the overall purple / violet sheen and pinkish-purple modification of ornamental spotting in *P. reticulata* by Pb. These cell types combine to produce not only background body coloration, but also increased reflectivity in the UV and/or near-UV spectrum.

*Poecilia reticulata* exist in a recently documented (Bias and Squire 2017a) polymorphic state; Autosomal Dominant Purple Body (*Pb/Pb and Pb/pb*) and non-Purple Body (*pb/pb*). The co-existence of the two phenotypes suggests a selective advantage under predation (crypsis) and in sexual selection (conspicuous pattern) under diverse ambient lighting conditions.

Pb has been identified as the first polymorphic autosomal gene to be described as existent in high frequencies in wild, feral and Domestic Guppy populations. It is capable of pleiotropic effects on all existing color and pattern elements at multiple loci. It should therefore be considered a strong candidate for further studies involving “the relationships between spectral and ultrastructure characteristics” in orange ornamentation, and extending to color and/or pattern as a whole as suggested by Kottler (2014). A mechanism is identified by which Pb is capable of balancing overall color and pattern polymorphisms, in turn providing fitness through heterozygosity in diverse complex habitats (Bias and Squire 2017a). We hope that Purple will be mapped to its linkage group.

## Photo Imaging

Photos by the senior author were taken with a Fujifilm FinePix HS25EXR; settings Macro, AF: center, Auto Focus: continuous, varying Exposure Compensation, Image Size 16:9, Image Quality: Fine, ISO: 200, Film Simulation: Astia/Soft, White Balance: 0, Tone: STD, Dynamic Range: 200, Sharpness: STD, Noise Reduction: High, Intelligent Sharpness: On. Lens: Fujinon 30x Optical Zoom. Flash: External mounted EF-42 Slave Flash; settings at EV: 0.0, 35mm, PR1/1, Flash: -2/3. Photos cropped or brightness adjusted when needed with Microsoft Office 2010 Picture Manager and Adobe Photoshop CS5. All photos by the senior author.

## Microscopy

All Digital Image processing by conventional bright and dark field equipment. AmScope M158C. Camera(s): 1. MD35, Resolution: 0.3MP 2. MD200, Resolution: 2MP USB Digital, Sensor: Aptina (Color), Sensor Type: CMOS. Software: AmScope for Windows. An attempt was made to restrict ambient light during both daytime and nighttime imaging of specimens. Imaging was performed with reflected or transmitted practical light sources as indicated. Where delineation in results warranted, a series of three photos from each location were taken and presented in the results; reflected (top light only), transmitted (bottom light only), combined reflected + transmitted (top and bottom light).

For purposes of this study low resolution photos were often preferred over higher resolution for clarity at settings of 40X, 100X or 400X. No images were stained. As identified, individual images are full body (non-dissected), or manually de-fleshed (dissected) skin samples. Samples were air dried for minimal time periods of less than one hour for aid in dissection.

All samples and images from right side of body, unless otherwise noted. No cover glass was utilized, to reduce damage to chromatophore shape, structure and positioning. No preservatives were used during imaging, though rehydration was done as needed for clarity. All photos were by the senior author.

## Ethics Statement

This study adhered to established ethical practices under AVMA Guidelines for the Euthanasia of Animals: 2013 Edition, S6.2.2 Physical Methods (6).

All euthanized specimens were photographed immediately, or as soon as possible, after temperature reduction (rapid chilling) in water (H20) at temperatures just above freezing (0°C) to avoid potential damage to tissue and chromatophores, while preserving maximum expression of motile xantho-erythrophores in Pb and non-Pb specimens. All anesthetized specimens were photographed immediately after short-term immersion in a mixture of 50% aged tank water (H20) and 50% carbonated water (H_2_CO_3_).

All dried specimens photographed immediately after rehydration in cold water (H_2_0). Prior euthanasia was by cold water (H_2_0) immersion at temperatures just above freezing (0 °C). MS-222 (Tricaine methanesulfonate) was not used to avoid the potential for reported damage and/or alterations to chromatophores, in particular melanophores, prior to slide preparation.

## Notes

This publication is number two (2) of four (4) by Bias and Squire in the study of Purple Body (*Pb*) in *Poecilia reticulata*:

1. The Cellular Expression and Genetics of an Established Polymorphism in *Poecilia reticulata*; “Purple Body, (*Pb*)” is an Autosomal Dominant Gene,
2. The Cellular Expression and Genetics of Purple Body (*Pb*) in *Poecilia reticulata*, and its Interactions with Asian Blau (*Ab*) and Blond (*bb*) under Reflected and Transmitted Light,
3. The Cellular Expression and Genetics of Purple Body (*Pb*) in the Ocular Media of the Guppy *Poecilia reticulata*,
4. The Phenotypic Expression of Purple Body (*Pb*) in Domestic Guppy Strains of *Poecilia reticulata*.

## Acknowledgements

To my best friend and wife Deana Bias, for her support and persistence over the last several years in this four part study… To my co-author and dear friend Rick Squire for his patience as a mentor… To those Domestic Breeders who willingly provided additionally needed pedigree strains and study populations for completion of this paper…

## Supporting Information

S1 Materials; Slide Specimen Photos

